# Neural mechanisms of credit assignment for inferred relationships in a structured world

**DOI:** 10.1101/2021.12.22.473879

**Authors:** Phillip P. Witkowski, Seongmin A. Park, Erie D. Boorman

**Affiliations:** Center for Mind and Brain, University of California Davis, Davis, CA, 95618; Department of Psychology, University of California Davis, Davis, CA, 95618

**Author notes:** Address correspondence to: Phillip P. Witkowski, Erie D. Boorman.

## Abstract

Animals have been proposed to abstract compact representations of a task’s structure that could, in principle, support accelerated learning and flexible behavior. Whether and how such abstracted representations may be used to assign credit for inferred, but unobserved, relationships in structured environments are unknown. Here, we develop a novel hierarchical reversal-learning task and Bayesian learning model to assess the computational and neural mechanisms underlying how humans infer specific choice-outcome associations via structured knowledge. We find that the medial prefrontal cortex (mPFC) efficiently represents hierarchically related choice-outcome associations governed by the same latent cause, using a generalized code to assign credit for both experienced and inferred outcomes. Furthermore, mPFC and lateral orbital frontal cortex track the inferred current “position” within a latent association space that generalizes over stimuli. Collectively, these findings demonstrate the importance both of tracking the current position in an abstracted task space and efficient, generalizable representations in prefrontal cortex for supporting flexible learning and inference in structured environments.

## Introduction

Much of human and animal behavior relies on the ability to effectively represent the environment and infer the true state of the world, which in turn supports effective decision making. For example, the value of taking a vacation depends not only on the weather in your current location but the weather in other locales, which are systematically related to your own. Observing cold winter weather in Chicago (northern hemisphere) predicts summer weather in the southern hemisphere, making a trip to Santiago, Chile, all the more valuable. In this situation, your brain needs both the ability to represent the underlying structure of the world (e.g., the inverse relationship between weather in each hemisphere) and the ability to assign credit for an inferred outcome (warm weather in Santiago) given an observed outcome (cold weather in Chicago). While this inference process is critical to flexible learning, the neural substrates that support credit assignment for inferred outcomes in real-world hierarchical environments are still unknown. In the current study, we test the hypothesis that the prefrontal cortex efficiently represents a hierarchical task space and uses this to infer unseen outcomes and assign credit to the appropriate latent cause.

Knowledge about the relational structure of environmental and task states is thought to be stored in representations called cognitive maps (Behrens et al., 2018; Gershman & Niv, 2010; O’Keefe & Nadel, 1978; Schuck, Cai, Wilson, Niv, et al., 2016; Tolman, 1948; Wilson et al., 2014). These representations contain information critical to goal-directed behavior, encoding relationships between positions or task states in an efficient manner. For example, outside of physical space, cognitive maps might contain relational knowledge about transition probabilities between states, choice-outcome contingencies, or how these contingencies change over time (Baram et al., 2021; Boorman et al., 2016; Daw et al., 2011; Hampton et al., 2006). In principle, cognitive maps are powerful because they allow for rapid updating when the state of the environment shifts (Bartolo & Averbeck, 2020; Boorman et al., 2021) and generalization to similarly structured tasks (Baram et al., 2021; Behrens et al., 2018; Franklin & Frank, 2018; Whittington et al., 2020). Within the prefrontal cortex, the lateral orbitofrontal cortex (lOFC) and medial prefrontal cortex (mPFC), in particular, have previously been implicated in using a model of the task’s structure, or an abstracted cognitive map of the task space, to assign credit for specific rewards to specific past choices or causes (Boorman et al., 2013, 2016; Jocham et al., 2016; Takahashi et al., 2011; Tanaka et al., 2008; Walton et al., 2010; Wilson et al., 2014). However, the neural mechanisms that underlie assigning credit to latent causes that generalize to inferred, but unseen, relationships in structured environments remain poorly understood.

To support credit assignment, prefrontal cortex may also play a critical role in tracking the state of knowledge within abstract task spaces. Unobservable task-relevant information that defines the current task state has been found during multi-step sequential tasks in OFC (Schuck, Cai, Wilson, Niv, et al., 2016; Wilson et al., 2014; Zhou et al., 2020). Moreover, recent work has pointed to interactions between the OFC and hippocampus that would allow the brain to track “positions” along trajectories through abstract task spaces to guide value-based decision making (Knudsen & Wallis, 2020; Zhou et al., 2019), with neurons in the anterior hippocampus coding the relative position along trajectories through the 3D abstract value space defined by each option’s current estimated value (Knudsen & Wallis, 2021). Recent advances in approaches to measure the neural representations of cognitive maps with functional magnetic resonance imaging (fMRI) have likewise identified abstracted cognitive maps of latent task spaces in human hippocampus and OFC (Clarke et al., 2019; Garvert et al., 2017; Park et al., 2020, 2021; Schapiro et al., 2016). Together, these insights suggest a new framework that may be extended to understanding associative learning in structured tasks: the brain might track the inferred position of hierarchically related associations in an abstracted “association space” that generalizes over choice stimuli for efficient model-based inferences and rapid updating.

In the current study, we address these questions using a “hierarchical reversal -learning task”, which required participants to use knowledge about hierarchical relationships to infer unobserved outcomes and make effective goal-directed decisions. We show that mPFC is a critical region both for efficiently representing choice-outcome relationships governed by a shared latent cause and for updating inferred choice-outcome associations at the time of feedback. Finally, we find that the lOFC and mPFC encode the inferred “position” within an abstracted association space for choice-outcome associations governed by the same latent cause.

## Results

### Hierarchical reversal-learning task

Participants completed a “hierarchical reversal-learning task” in which they tracked the probability that each of four fractal shapes would lead to either of two gift cards for one of two different online stores (Fig.1A). On each trial, participants choose between two of the four shapes based on two pieces of information: estimates of the probability that a particular shape will lead to a particular outcome, and the randomly generated potential payout indicated for each outcome (Fig.1B). Importantly, the set of fractal shapes were organized hierarchically into two independent systems of inverse pairs. Shapes A and B formed “System 1”, while shapes C and D formed “System 2”. This hierarchical organization gave participants the opportunity to infer unobserved outcomes for an unchosen shape when observing the outcomes derived from choosing the system pair. For example, participants could track the probability that A leads to outcome 1, by observing the frequency that B leads to outcome 2. Because the two systems were independent of each other, however, nothing could be learned about shapes C or D from observing the outcomes of shapes A or B. Participants completed a total of 160 trials across two sessions, during which the associative contingencies reversed three times (Fig.1C). Participants were told that one trial would be selected at random to count “for real” at the end of the experiment and they would be given money proportional to the number of points won on the gift card they received for that trial.

**Fig.1.**
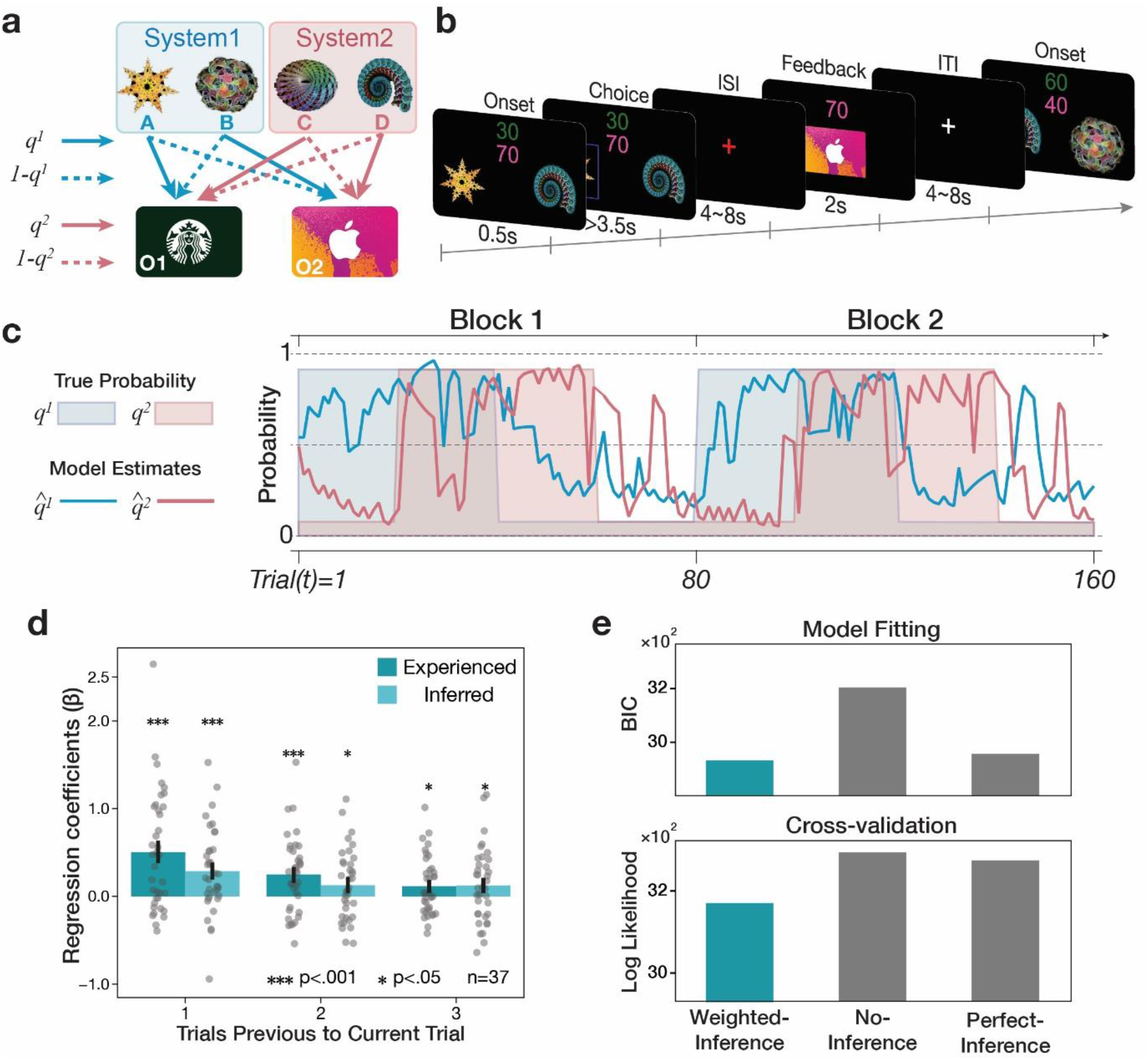
Learning Task Design and Behavioral Results. **a**) Four fractal shapes were organized hierarchically into two independent systems of inversely related pairs. This meant that participants could infer the outcome of one object (e.g., shape B) after observing the outcome from choosing its system pair (e.g., shape A). **b**) Illustration of the fMRI task. Participants were presented with 2 of the 4 shapes to choose from in each trial. They chose between the shapes on the basis of two pieces of information: their estimate of the transition probabilities ( *q*^1^, *q*^2^) that an object would lead to either gift card outcome, and the randomly generated number of points they could potentially win on each gift card if obtained. The color of each number indicated the identity of the outcome on which that number of points could be won. In the example, green indicates the number of points for the Starbucks gift, while pink indicated the number of points for iTunes. Next, they observed the outcome of their choice (the gift card and amount) after a delay. **c**) Example of a participant’s learning trajectory as the task unfolded. Shaded regions indicated the true associations for system 1 (*q*^1^, blue) and system 2 (*q*^2^, red). Each system reversed 3 times during the experiment, switching *q*^1^ and *q*^2^ to 1-*q*^1^ and 1-*q*^2^, respectively. Blue and Red lines indicate the estimated values of 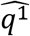 and 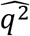 based on the weighted-inference learning model (see computational models for details). **d**) Results of a logistic regression analysis which shows the influence of past choices and outcomes on the current choice. Both experienced and inferred past choice-outcome associations significantly predicted current choice. As expected, this influence decreased for trials further in the past. Height of the bars represents the mean of regression coefficients ± SEM. **e**) Results of model comparisons using BIC (top) and 8-fold cross-validation (bottom) for weighted-inference, no-inference, and perfect-inference models (see computational models for details)

### Behavioral results

Optimal behavior in this task required that participants tracked which stimulus choices led to which of the two outcomes and used that knowledge to make decisions on the current trial. We characterized the influence of previous choice outcomes using logistic regression models that predicted the odds of choosing a certain shape given the currently desired outcome (i.e., the stimulus with a higher payoff) and outcomes resulting from the last three times that shape was chosen (EQ.8). Note that available choice stimuli changed on each trial so these outcomes may be more than 3 consecutive trials into the past. Critically, we also included the outcomes that could be inferred from choosing the system pair – the source of inferred information in our task – in the regression model. If participants utilized both experienced and inferred outcomes to learn, reinforcement learning theory predicts positive effects of each type of outcome that decline exponentially over time into the past (Bayer & Glimcher, 2005; Sugrue et al., 2005).

This analysis showed significant effects for all three experienced and inferred choice-outcome pairs going three choices into the past (all t(36)’s > 1.94, all p’s < .05) (Fig.1D). This learning-model agnostic analysis confirms that subjects learned from both the experienced and inferred choice-outcomes associations and utilized this information to make decisions on the current trial. We compared the magnitude of regression coefficients between experienced and inferred outcomes over time using a two-factor ANOVA. We found an expected main effect of time (F(2,72)= 5.63, p<.01), showing that outcomes from trials further in the past were less influential on the current choice. However, the magnitude of effects from experienced outcomes were not found to be significantly greater than those from inferred trials (F(1,36)=2.97, p=.09), and there was no significant interaction between outcome type and time (F(2,72)= 2.34, p=.10). Finally, the analysis showed no effect of the previous outcome’s reward magnitude on the subsequent trial’s choice (t(36)=-1.03, p=.85), consistent with the fact that they were generated randomly on each trial and there was no advantage to tracking rewards between trials in our task. Taken together, this analysis shows that subjects learned from both experienced and inferred outcomes and that directly experienced outcomes did not have a significant advantage in guiding future decisions relative to inferred outcomes in our task (results were similar when incorporating the subjective value of each outcome into the analysis (Fig.S1)).

To estimate subjects’ trial-by-trial beliefs about stimulus-outcome associations, we fit each participant’s choices to a Bayesian reversal learning model (see methods) that utilized the history of outcomes observed from their choices, and outcomes inferred from the system pair. The best-fitting “weighted inference model” jointly estimates the stimulus-outcome (transition) probability and the reversal probability and included three free parameters: *α*, an indifference term capturing the subjective preference for one outcome over the other; *β*, an inverse temperature term capturing participants’ sensitivity to differences in choice values; and *γ*, an inference weight term which weighted the posterior belief in choice associations for experienced relative to inferred outcomes, reflecting the amount of information each subject derived from a directly experienced outcome relative to an inferred outcome (see EQs 2–6 and Table S1 for the distribution of parameter estimates).

We compared the “weighted inference model” to two alternatives which did not include *γ*, but instead assumed the participants learned nothing from inference (“no-inference model”) or learned perfectly from inferred and experienced information (“perfect-inference model”), using Bayesian Information Criterion (BIC) (EQ.7). The weighted inference model was found to best capture choice data across subjects compared to these alternative models (lowest summed BIC across subjects), showing that that the weighted inference model (BIC=7266.34) captured meaningful differences in participants’ ability to infer from unobserved data (no-inference model BIC=7401.76; perfect-inference model BIC=(7278.12) (Fig.1E). We further confirmed this finding using forward chaining cross validation (k=8; Bergmeir & Benítez, 2012) to show that this model predicted out-of-sample choices better than models that assumed either no inference or perfect inference.

Finally, we tested if subjects’ choices were a sigmoidal function of the estimated expected value of each choice option using the weighted inference model (likelihood ratio test (LRT) = 60.44, p=7.32×10^−17^). Fig.S1 shows the highly significant results of a multilevel logistic regression model predicting the subjects’ choices given the expected value (EQ.4) difference between the two options on each trial.

### Neural substrates of belief updating from experienced and inferred outcomes

Our next analysis sought to identify the network of brain regions that support updating of choice-outcome associations by combining information from experienced and inferred outcomes at the time of feedback. We defined the belief update from feedback as the Kullback-Leibler divergence (D_KL_) between prior and the posterior beliefs after observing the outcome on each trial, also called the “Bayesian surprise” (Iglesias et al., 2013; Schwartenbeck et al., 2016a). Because participants may learn through both experienced and inferred outcomes, the total update on a given trial is the sum of the D_KL_ for experienced and inferred choice-outcome associations (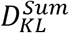; EQ.12). We used 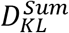 as a parametric modulator of blood oxygen-level-dependent (BOLD) activity during feedback (see GLM 1) and found clusters of positive effects in pre-supplementary motor area/dorsal anterior cingulate (preSMA/dACC) (peak voxel, [x,y,z]=[0, 18, 50], t(36)=7.31), bilateral DLPFC (right, [x,y,z]=[46,24,48], t(36)=5.90; left, [x,y,z]=[ 36,8,36], t(36)=6.22) and bilateral anterior insula (right, [x,y,z]=[32, 26, 0], t(36)=5.35; left, [x,y,z]=[−32, 22, 2], t(36)=5.69), (all whole-brain cluster-corrected with permutation-based threshold-free cluster enhancement (TFCE) (Smith & Nichols, 2009) at p_TFCE_ <.05), suggesting these regions encode updates to the system of choice-outcome associations (Fig.S2, Table S2; see Fig.S3 for reward prediction error effects).

Next, we tested for regions that carried additional information about updating derived from inferred information. We did this by calculating the D_KL_ for the “no inference” model 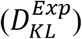, which quantified the update on the current trial if no inference occurred (i.e., only experienced information was used in the update). We then used the “weighted-inference model” to compute the D_KL_ given the subject-specific weighting of inferred information 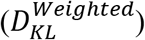. We computed the difference between these regressors (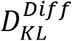, *EQ*. 12) to quantify the additional updating that occurs when inferred information is combined with directly experienced information to update beliefs. We used the trial-by-trial estimates of 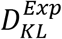, and 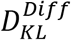 as parametric modulators of BOLD activity at the time of feedback (see GLM 2), to identify regions that reflected the additional update gained from inference, even while controlling for updates due to experienced outcomes only. We found significant positive effects of 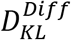 in clusters in preSMA/dACC ( [x,y,z]=[4, 20, 48], t(36)=5.56), bilateral DLPFC (right, [x,y,z]=[40, 38, 18], t(36)=4.37; left, [x,y,z]=[−44, 6, −24], t(36)=5.24) and bilateral anterior insula (right, [x,y,z]=[32, 20, 4], t(36)=5.04; left, [x,y,z]=[−30, 22, −1], t(36)=5.36) (Fig.2A, Table S3). These results implicate this network in supporting the additional updating of beliefs about transition probabilities from inferred outcomes at feedback.

**Fig.2.**
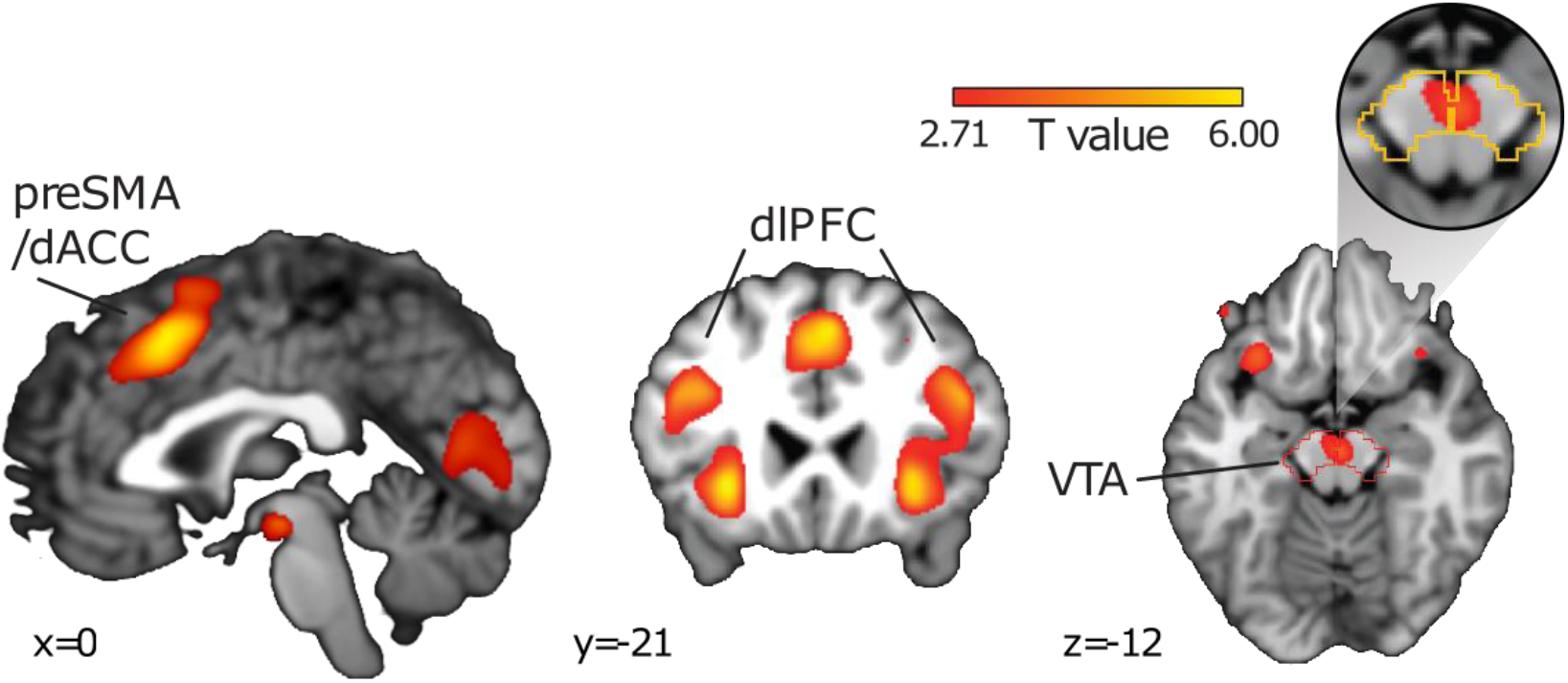
Network of Regions that Reflect Additional Update for Inferred Information. Sagittal and coronal slices through t-statistic maps display brain regions whose activity at feedback reflected the additional information gained from including inferred information compared to only experienced information 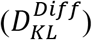. For illustration, maps display regions at a threshold of t(36)=2.71, p<.005, uncorrected.

Recent studies have suggested that activity in the dopaminergic midbrain encodes prediction errors not only about reward value but also about outcome identity or ‘task state’ (Boorman et al., 2016; Gershman & Uchida, 2019; Howard & Kahnt, 2018; Iglesias et al., 2013; Langdon et al., 2018; Sharpe et al., 2017; Suarez et al., 2019). As such, we tested whether activity in the dopaminergic midbrain, in particular the ventral tegmental area (VTA), would also reflect the additional update of transition probabilities based on inferred information 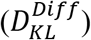, using an independently defined region-of-interest (ROI) over the VTA and substantia nigra (SN) (Diaconescu et al., 2017). Consistent with our prediction, we found a significant positive effect of the combined update at the time feedback was delivered in the VTA ([x,y,z]=[2, 18, −12], t(36)=3.39, p_TFCE_ <.05, ROI corrected), independent of reward prediction error. Notably, we found no significant effect of the reward prediction error (EQ.6) in the same VTA/SN ROI (Fig.S3), consistent with the fact that there was no incentive to learn from reward magnitudes in our task, and subjects did not show a behavioral effect of *learning* from reward magnitudes, as shown above. Collectively, this suggests that the VTA BOLD signal aligns with the instrumentally relevant variable to track in our task, and, importantly, incorporates inferred information based on knowledge of the task structure (Fig.2B).

### mPFC represents latent causes and assigns credit to inferred outcomes

We hypothesized that the brain would reinstate the latent cause using an efficient code that generalizes over stimuli and outcomes governed by the same cause at feedback time. If participants retrieve representations of structural relationships at feedback to appropriately assign credit to the latent association, we would expect to decode the representations associated with the common causes that arise in trials where the systems’ pairs led to opposite outcomes. To probe which brain regions assigned credit to a shared representation for shapes governed by the same causal relationship (i.e., shapes part of the same system), we performed a multivariate pattern analysis (MVPA) on activity patterns at feedback, the critical time for credit assignment. First, we trained pattern-based classifiers (linear support-vector machines) to classify the chosen stimulus and its associated outcome identity at the time of feedback (e.g., A→O1), and then used the resulting feature weights to decode from patterns of activation on trials where the system pair led to the opposite outcome through the same causal relationship (e.g., B→O2) (Fig.3A, see supplement for details on decoding procedure). Importantly, this analysis controlled for both the shape stimulus and outcome identity such that no sensory information, neither the previous choice stimulus nor reward outcome identity, was shared between training and test sets. Thus, decoding is only possible if these events share information about the same causal relationships that bind shapes in the same system.

**Fig.3.**
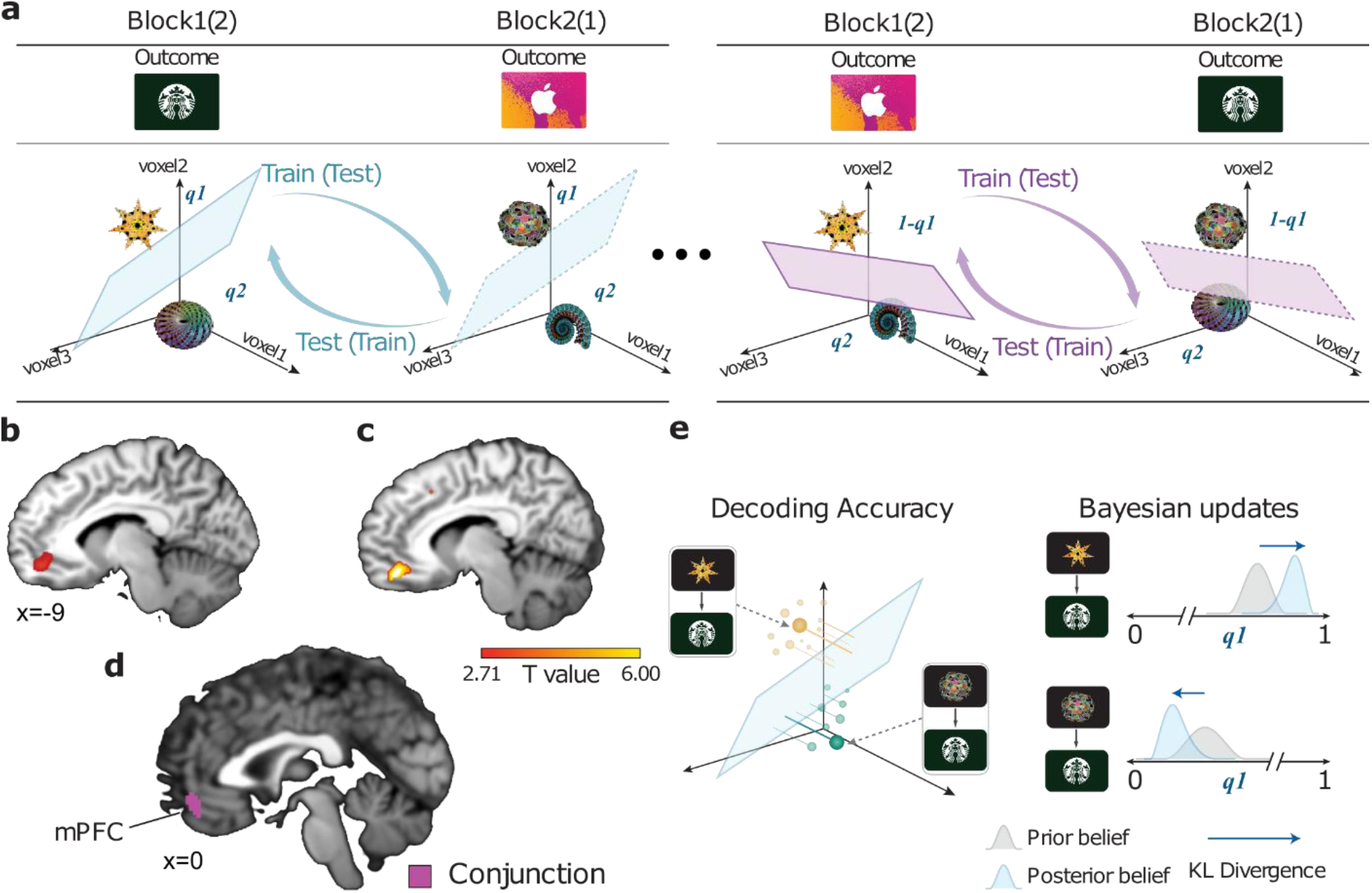
Medial PFC Carries Representations of the Latent Cause to Assign Credit to Inferred Outcomes. **a**) Illustration of the decoding procedure used to decode the latent cause. We first trained a linear SVM on specific shape-outcome combinations from each system (e.g., A→O1 and C→O1) then used it to classify the system pairs which led to the opposite outcome (B→O2 and D→O2). No information other than the latent cause was shared between training and testing trials. In a separate analysis (**e**), we correlated the amount of information about the latent cause in each trial (distance from SVM hyperplane) with the magnitude of updates estimated by the weighted-inference learning model (see multivariate analysis for details). **b**) Sagittal slice through t-statistic map showing effects of decoding of the latent cause from analysis depicted in **a** in mPFC (SVC within an *a priori* mPFC ROI), displayed using the same conventions as Fig.2. **c**) Same as **b** but shows regions where the magnitude of information decoded about the latent cause was significantly correlated with 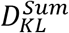 (SVC in mPFC ROI). **d**) Conjunction t-statistic map showing overlapping regions of **b)** and **c)** (p<.05 uncorrected).

We began by conducting a whole-brain searchlight analysis to estimate decoding accuracy at each voxel in the brain (Kriegeskorte et al., 2008). Based on our *a priori* hypotheses concerning lOFC and mPFC in credit assignment (Boorman et al., 2013, 2016; Jocham et al., 2016; Tanaka et al., 2008; Walton et al., 2010), we tested anatomically defined ROIs (Glasser et al., 2016) of mPFC and lOFC that were hypothesized to contain these representations and used TFCE (Smith & Nichols, 2009) to correct for multiple comparisons. This analysis identified a significant cluster of voxels in the anterior portion of left mPFC ([x,y,z]=[−6,50,-10], t(36)=3.54, p_TFCE_ <.05 ROI corrected); Fig. 3B, Table S4). However, we found no significant clusters in lOFC bilaterally (all p>.05 uncorrected).

To more directly test whether these representations of the latent cause in mPFC relate to credit assignment during inference, we correlated the strength of representations of the latent cause in mPFC at the time of feedback with model-derived estimates of the updates to outcome contingencies within each system. We used the same SVM classifier to compute the decodability of system representations at feedback during each trial. We quantified the decodability of each representation as its distance to the SVM hyperplane (Schuck & Niv, 2019) and signed the distances such that correct classifications were positive and incorrect classifications were negative. As before, we defined the total trial-by-trial belief update as the 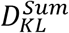 between the prior and posterior beliefs after having observed an outcome. This whole-brain analysis revealed a significant cluster in mPFC (Spearman rank correlation; [x,y,z]=[8,46,-10], t(36)=4.19; p_TFCE_ <.05 ROI corrected Fig. 3C, Table S5), which overlapped with the main effect of latent cause decoding (Fig. 3D; using the conjunction analysis with minimum statistics, at p <.05 uncorrected compared to conjunction null; (Nichols et al., 2005)). This finding shows enhanced representation of the common causal relationship with greater updating for credit assignment for both experienced and inferred outcomes at the time of feedback.

### lOFC and mPFC track inferred positions in a latent association space during learning

Our results have shown that mPFC contains a representation of underlying causal relationships that are used to infer information about related stimuli during feedback. Based on recent evidence showing that hippocampus and OFC may track the current position within a value or task space (Knudsen & Wallis, 2020, 2021; Park et al., 2020; Schuck, Cai, Wilson, Niv, et al., 2016), we hypothesized these regions may track the “position” of subjects’ current beliefs within an abstract “association space” for each system. To test this hypothesis, we used representational similarity analysis (RSA) to identify regions of the brain that coded relative “positions” within the inferred association space. That is, we sought to identify brain regions that had increasingly similar representations when subjects had increasingly similar beliefs about the choice-outcome contingencies for each system. We generated a model Representational Dissimilarity Matrix (RDM) that calculated the divergence (Jensen-Shannon Divergence (*D_JS_*); a symmetric measure of the distance between distributions, EQ.13) between model estimates of the posterior belief distributions about stimulus-outcome associations in a system (e.g., *q*^1^) computed from our weighted-inference learning model in each trial across sessions. We also generated a RDM of neural similarity from activity patterns measured within a searchlight during the inter-trial interval (ITI) by calculating the Euclidean distance between voxel patterns in each trial across sessions. We hypothesized that regions tracking one’s current position in the association space would show increasingly greater representational similarity for trials that had increasingly similar posterior beliefs about the specific position of a configuration of associations within a system. We reasoned that if subjects were tracking the latent cause governing a system of associations (A→O1, B → O2), then this coding should be independent of the specific choice made within that system (e.g., include both A (C) and B (D) choices for system 1 (2)) (see Fig. 4A and Methods).

**Fig.4.**
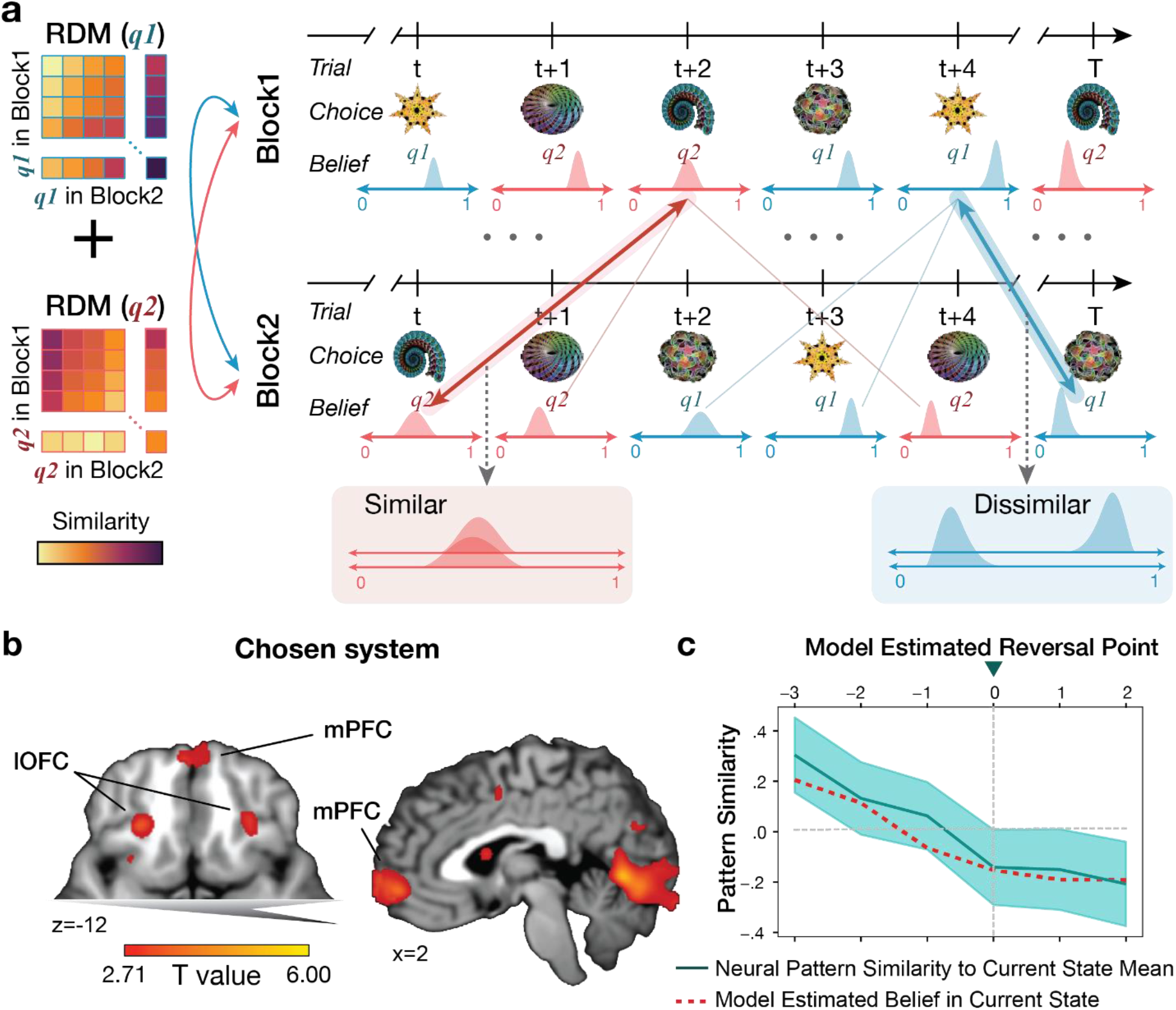
Lateral OFC and Medial PFC Track Inferred Positions within Latent Association Space During Learning. **a**) Conceptual illustration of the RSA procedure used to test for estimated position within the latent association space. We constructed a model RDM that measured the dissimilarity of posterior beliefs (*D_JS_*), estimated by the weighted inference learner, across trials in separate blocks. Only trials in which shapes from the same system were chosen by the participant were compared across blocks. Separate RDMs for each system were then compared to neural RDMs computed from the ITI period of the same trials, using the Euclidean distance between voxel activation patterns on these trials from different blocks as the measure of dissimilarity. Model and neural RDMs were then compared using linear regression (see “Representational Similarity analysis of Association Space” for details). **b**) Axial and sagittal slices through t-statistic map displaying regions in which the model RDM was significantly related to the neural RDM. Maps are displayed with the same conventions as in Fig.2. The clusters survived small volume correction within an *a priori* defined lOFC ROI (axial slice) and mPFC ROI (sagittal slice). **c**) Visualization of the relationship between model-estimated reversal points and neural pattern similarity. Dashed vertical line indicates a reversal point, where 0 is the trial directly after a reversal in the configuration of each system, as estimated by the weighted-inference learner. Green line represents the neural similarity of the activation patterns in lOFC on each trial immediately preceding and subsequent to the reversal point, compared to a “template pattern” – defined as the average pattern from trials with the same configuration as those prior to the reversal point, but from the other block. Red line shows the model-derived belief estimate on the same trials. Note the corresponding shift in the model estimate and neural data from pre- to post-reversal.

We tested this hypothesis by constructing a general linear model that predicted values of the neural RDM while controlling for other possible explanations of neural similarity, using the *D_JS_* model RDM along with 5 control RDMs. These alternative RDMs controlled for the effect of the position in association space of the unchosen system for the current trial as well as similarity of the recently observed outcome, identity of the chosen object, identity of the unchosen object, magnitude of the RPE, and physical response made (see methods). We focus on the ITI following recent evidence of positional coding in an abstract value space during the ITI in monkey hippocampal single unit recording (Knudsen and Wallis, 2021). The RDM representing task “position” revealed significant effects in a network of regions including bilateral lateral OFC (left lOFC, [x,y,z]=[−26, 30, −12], t(36)=4.24, p_TFCE_ <.05 ROI corrected; right lOFC, [x,y,z]=[28, 28, −14], t(36)=3.79, p_TFCE_ <.05 ROI corrected) and rostral mPFC ([x,y,z]=[4, 54, −4], t(36)=4.56, p_TFCE_ <.05 ROI corrected) (Fig.4A, Table S6). Indeed, visualization of pattern similarity in lOFC on the trials immediately before and after an inferred reversal point support this finding by revealing a shift in representation from the previous to the current belief state, in tandem with the shift in model estimates (Fig. 4C). This visualization showed positive pattern similarity to the current state prior to the reversal and shift to negative pattern similarity at the inferred reversal point. Collectively, these findings show that the lOFC and rostral mPFC track the current position in an abstract association space that generalizes over choices in the same system.

## Discussion

Understanding how the brain uses abstracted internal models to learn from unobserved, but inferred, outcomes is essential for understanding flexible behavior in complex environments. The current experiment adds to a growing body of work showing that the mPFC is critical to maintaining compact and generalizable representations of task-relevant variables (Baram et al., 2021; Behrens et al., 2018; Constantinescu et al., 2016; Iordanova et al., 2007; Morton et al., 2020; Samborska et al., 2021) but goes further to show these representations support credit assignment when outcomes can be inferred through shared hierarchical relationships. Our results show that mPFC selectively encodes the shared causal relationship between hierarchically related choice-outcome associations with a compact representation and leverages this code to assign credit for unseen, but inferred, choice-outcome associations. We also show that mPFC and lOFC code the current inferred position of the hierarchically related system within a common “association space” for each system, suggesting that these regions are integral for tracking the learner’s “position” within a latent association space as learning unfolds.

We designed a novel hierarchical reversal learning task to test the hypothesis that assigning credit for inferred outcomes depends on the reinstatement of a generalizable neural representation that links both experienced and inferred causal relationships (Liu et al., 2021). Prior evidence across species has implicated both the lOFC and the mPFC in credit assignment (Boorman et al., 2013; Chan et al., 2016; Jocham et al., 2016; Takahashi et al., 2011; Tanaka et al., 2008; Tsujimoto et al., 2009; Walton et al., 2010) but the precise functional roles attributed to each region remained unclear. Consistent with studies showing that mPFC contains condensed, low-dimension codes for structurally related items in the environment (Constantinescu et al., 2016; Doeller et al., 2010; Morton et al., 2020; Park et al., 2021; Samborska et al., 2021), we found that mPFC, but not OFC, reinstated the shared latent cause that governed two sets of stimulus-outcome associations in the same system. Importantly, this effect could not be explained by either the outcome’s identity or the identity of the chosen stimulus alone. Further, we show that the decodability of these representations in mPFC increases when subjects updated their estimates to a greater extent, which is consistent with prior work showing that representations in the mPFC are important for rapid updating between states (Klein-Flügge et al., 2019; Muller et al., 2019). These results suggest that generalized representations in mPFC are used for credit assignment at feedback, directly linking knowledge about causal structure to inference about unobserved outcomes. Moreover, they provide novel evidence that cognitive maps may be used to generate inferences about an untaken choice based on knowledge about the underlying relational task structure.

Our study also extends our understanding of the network of regions involved in updating choice-outcome associations, by showing that these regions also support updating from inferred outcomes using a model of the task’s hierarchical structure. A network of regions’ activity reflected the full learning update (D_KL_) from an outcome, including the VTA, pre-SMA/dACC, dorsolateral prefrontal cortex, ventrolateral prefrontal cortex/lOFC, and anterior insula, consistent with past studies investigating directly experienced outcomes/stimuli (Boorman et al., 2016; Iglesias et al., 2013; Schwartenbeck et al., 2016b). These findings support the view that dopaminergic precision-weighted prediction errors modulate both local cortical and long-distance cortico-cortical and cortico-striatal synapses within a similar network of regions during incremental learning (Stephan et al., 2015). Notably, dopaminergic neurons in the VTA are known to signal reward prediction errors (Bayer & Glimcher, 2005; Montague et al., 1996; Schultz et al., 1997) but more recent work has suggested this role extends to updating value-neutral associations between states or outcome identities. Indeed, activity in the VTA is modulated by errors in predicted outcome identity (Howard & Kahnt, 2018; Iglesias et al., 2013, 2021; Oemisch et al., 2019; Suarez et al., 2019; Takahashi et al., 2017) and belief updating about the state of associative relationships in the environment (Schwartenbeck et al., 2016a; Sharpe et al., 2017) which have been shown to play a causal role in learning such value-neutral associations (Langdon et al., 2018; Sharpe et al., 2017). Here, we show activity in the VTA quantitatively encodes precision-weighted prediction errors about the state of hierarchically related choice-outcome associations, integrating information from both experienced and inferred outcomes. Furthermore, this signal only reflected how much to learn about the instrumentally relevant variable and did not track learning-irrelevant, but nonetheless rewarding, outcomes. We found no evidence that the VTA signal incorporated the monetary reward value obtained at feedback, which in our task is irrelevant for future behavior. This is consistent with the absence of any effect of reward magnitude on learning behaviorally. Taken together, our findings highlight the importance of dopamine in updating model-based associations through inference.

Finally, we show that a network of brain regions including lOFC and mPFC track the inferred position in a latent association space that generalizes over choice-outcome associations within a system. We found that lOFC and rostral mPFC showed relational coding corresponding to the position in the hierarchically related choice-outcome association space, such that activation patterns were increasingly similar when the expectation and precision of beliefs about associations within a system were more similar. This finding dovetails with recent studies showing that relational position in a wide range of abstract spaces are coded by medial temporal lobe and orbitofrontal cortex (Constantinescu et al., 2016; Knudsen & Wallis, 2021; Park et al., 2020; Theves et al., 2019). Here, we show this coding scheme applies to a hierarchically general latent causal space in lOFC and mPFC that reflects both the certainty and confidence in learned choice-outcome associations (Pouget et al., 2016). While we did not find any significant effects in hippocampus at the thresholds used, there was a subthreshold correlation in the head of the right hippocampus (p_TFCE_ <.08 ROI corrected, Fig.S4). Recent pioneering studies using closed-loop theta stimulation in monkeys have identified a causal role for hippocampal input to a homologous region of lOFC (Brodmann Area 13) during the ITI of a reward-guided learning task (Knudsen & Wallis, 2020). A second study elaborated these findings by showing that hippocampal neurons coded for direction dependent “positions” in the monkeys’ trajectory through an abstract 3D value space (Knudsen & Wallis, 2021). Taken together with our findings, this suggests that representations of learning trajectories in lOFC and mPFC may be derived from hippocampal relational codes, which are input to these regions through direct anatomical connections (Barbas & Blatt, 1995). In our study, these codes can be used for accurate credit assignment and inference. More generally, our findings support the theory that the OFC represents an animal’s current position in a task space when its position cannot be directly observed (Schuck, Cai, Wilson, & Niv, 2016; Stalnaker et al., 2015; Wilson et al., 2014; Zhou et al., 2020).

An intriguing open question is whether lOFC would reactivate specific individual past choices, as opposed to generalizable latent causes with a common code, for credit assignment to specific past choices. Previous work has shown that OFC reactivates choices which led to the currently observed outcome specifically at outcome time (Tsujimoto et al., 2009), and may trigger reactivation of sensory representations via descending anatomical connections between areas of posterior and lateral OFC and several sensory cortical regions (Carmichael & Price, 1995; Cavada et al., 2000). Whether or not the same mechanism underlies credit assignment for inferred stimuli is unknown. Notably, we did not find any significant decoding of the chosen stimulus identity alone at feedback anywhere in the brain at our threshold used (p_TFCE_ <.05). This finding is consistent with our fMRI decoding and behavioral analyses showing that by and large subjects treated stimulus-outcome associations governed by the same cause as a unitary representation, rather than treating its individual associations distinctly. Future work can elaborate these mechanisms by testing whether the appropriate inferred choices are reactivated in a modality-specific sensory cortex during learning.

In conclusion, we find that the human brain represents latent causes with compact representations in mPFC, which support updating during credit assignment to inferred relationships. Further, relational codes in both lOFC and mPFC track learning positions along trajectories within an abstract association space that generalizes over stimuli, and rapidly update the actor’s position as learning dynamically unfolds. Collectively, these findings support a novel framework for understanding how the human brain learns in hierarchically structured settings that abound in the real world.

## Methods

### Subjects

Forty subjects (25 females; mean age = 20.5) were recruited from the general population around University of California, Davis. None of the participants reported a history of neurological or psychiatric disorders. Subjects either received either course credit or money ($15/hour) for participation in the experiment. Two subjects were removed due to excessive motion during scanning (head movement > 3mm), while a third subject was removed for excessive dropout in ventral regions of the prefrontal cortex that are of interest to this study. Thus, the final sample included 37 subjects (22 Females; mean age = 20.5). All procedures were approved by the University of California, Davis IRB. Participants gave written consent before the experiment.

### Hierarchical-reversal-learning-task

#### Task instruction

Subjects completed a “hierarchical-reversal-learning-task” in which they tracked associations between abstract shapes (choices) and reward identities (outcomes) to optimize the possibility of larger rewards at the end of the experiment (Fig.1A). On each trial, subjects were presented with 2 of 4 different fractal shapes from which to choose. Two numbers between 0 and 100 were presented at the top of the screen in unique colors. The color of the numbers corresponded to the identity of the gift-cards that the subject could win, and the magnitudes corresponded to the point value of the reward on the current trial. For example, a pink “42” meant that subjects could win 42 points on an iTunes gift-card while a green “58” meant they could win 58 points on a Starbucks gift card. The cumulative number of points available on each trial was always equal to 100. Subjects were told that the point values were randomly chosen on each trial and there was no point to tracking them.

Each shape had a certain probability of leading to one outcome and the inverse probability of leading to the other. For example, at the start of the experiment shape “A” would lead to the Starbucks gift-card with probability *q*^1^ and the iTunes gift-card with probability 1-*q*^1^. However, these true probabilities would reverse such that a given shape would lead to each outcome with opposite probabilities. Continuing with our example, after a reversal, shape “A” would lead to an iTunes gift card with probability *q*^1^ probability and a Starbucks gift card with 1-*q*^1^ probability. The point values (reward magnitudes) for each outcome were generated randomly from the range 0-100 on each trial, meaning that subjects did not need to track the reward magnitudes between trials. Instead, to maximize rewards, participants had to track the probability a shape would lead to each of the outcomes over trials and combine this with the reward magnitudes associated with each outcome on the current trial to guide their decisions based on their subjective preference.

Crucially, the shapes were organized such that they formed 2 sets of inversely related “systems”. Shapes within a system always led to opposite outcomes and had inverted outcome probabilities. Shapes A and B were paired (system 1) and shapes C and D were paired (system 2). The inverse relationships within a system allowed subjects to learn the probability that a shape would lead to a specific outcome by observing the choice-outcome relationship of the other shape within the same pair. For example, experiencing that shape A led to Starbucks would also give you the knowledge that if shape B were available and it was chosen, the outcome would have been iTunes. The same relationship was true for shapes C and D. Between systems, observations were completely independent of each other such that observing an outcome from choosing A or B gives no information about the likely outcomes of choosing shapes C or D. These structural relationships between choice options and outcomes within a system, and the independence of items between systems, was clearly explained to participants before the experiment began.

However, subjects did not have any prior knowledge about choice-outcome associations, and when reversals in choice-outcome associations occurred, or how many times reversals would occur (three times for each system, see Fig.1A). Therefore, subjects needed to infer both associative contingency for each choice and when reversals had occurred from their choices and outcome histories during experiments.

#### Stimuli

Four visually distinct unfamiliar fractal images were chosen such that the visual similarity between any two items were minimal and were presented to all participants as choice options. Images for system 1 and those for system 2 were randomized across participants.

Two types of reward identities (two gift-cards images) were chosen from 7 different gift-cards from stores familiar to participants: Best-Buy (blue), Barnes and Noble (tan), iTunes (pink), Regal (purple), REI (orange), Sephora (white), and Starbucks (green). The two reward identities were chosen prior to the fMRI experiment based on participant’s preference ratings. Subjects rated their preference level for each of these gift cards presented in a random order on a 1-100 scale. A pair of gift-cards having the minimum difference among four most highly preferred were selected per individual participant. These two gift-cards were assigned to outcome 1 (O1) and outcome 2 (O2), counterbalanced across subjects, and presented during fMRI experiment. This procedure allowed us to minimize potential biases from initial preferences in choices during the reversal learning task, while maintaining a high desirability for each outcome. All stimuli in each phase were presented on a computer running Psychopy v1.84 (Peirce, 2009).

#### Task-Schedule and Procedure

We generated two separate schedules that determined which choice options (shapes) would be presented on each trial and when reversals would occur. In this experiment, there were six possible unique combinations of four choice stimulus on any trial. In the experiment schedule, none of the same combination was repeated twice in consecutive trials. Further, we optimized the schedule such that an ideal Bayesian learner (perfect inference model; see Computational models) would choose each shape and receive each outcome approximately equally, given an equal preference between outcome identities. This was important because it minimized the potential for sampling bias in planned multivariate analyses (see Multivariate Analyses). Each schedule had predetermined reversal points where the choice-outcome associations switched (e.g., *q*^1^ →1-*q*^1^ and 1-*q*^1^→*q*^1^) for a given system. During fMRI experiments system 1 reversed every 40 trials starting from the first trial onwards, while system 2 reversed every 40 trials starting from the 20^th^ trial onwards, making the state of each system independent of each other. The independent reversal points of two systems made it so participants were not able to learn the choice-outcome associations of one system from that of the other.

Subjects completed two blocks of 80 trials (160 trials total). Before the fMRI experiment, subjects were instructed that one trial would be chosen at random to count “for real” and would be used to calculate the subjects reward for the experiment. This makes each choice independent. Therefore, participants need to make an optimal decision for every trial to maximize their rewards. At the end of experiment, we randomly selected one trial and gave a reward proportionate to the number of points earned on the specific gift card received on that trial. The minimum reward given was $5 while the maximum value was $25.

#### Behavioral Training

To familiarize subjects with the task, all subjects completed a behavioral training session before the fMRI experiment. After behavioral training participants performed the fMRI experiment on different day within a week. The task used for behavioral training was the same with the fMRI task except for slight modifications to aid learning. During behavioral training the experimenter guided subjects through the first 30 practice choice trials to ensure that subjects understood the task, then left participants to complete the rest of the trials on their own. In addition, to ensure that subjects tracked the relationship between paired stimuli, subjects were tested every 10th trial on the relationship between shapes, by asking them to connect shapes in the same pair with a single line (Fig.S5). The subjects received feedback via the line color - an incorrect pairing resulted in the line turning red, while a correct pairing turned the line green. During behavioral training participants learned the task with the same fractal images assigned to the same systems. However, we used 2 faux outcome identities (Zappos and Netflix) that would not be available for rewards during the fMRI experiment. Participants who understood the task well and performed well (model fit negative log-likelihood <130) were invited to return for fMRI experiments. Among 48 participants who initially enrolled the experiment, 40 participants participated in fMRI experiment.

### Computational models

#### Weighted-Inference learning model

We designed a Bayesian computational model to predict the choice of participants in each trial *t* based on one’s choice and outcome history and available choice options and reward magnitudes of the current trial. On each trial, the model estimated the contingency that choosing a given shape (*S*) would lead to outcome 1 (*O1*), and by definition led to outcome 2 (*O2*) with the inverse probability which is denoted as follows:

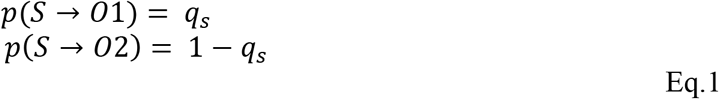

Choice-outcome contingencies for all shapes were modeled as separate distributions, but beliefs about contingencies for shapes in the same system were related through an inference term (*γ* where 0 < *γ* < ∞), which takes account to what extent an individual participant learns and updates *q_s_* from direct experiences (the outcome *y* after choosing *S*) compared to that from inferred outcomes (the outcome *y′* if you had chosen *S′* where, *S* and *S′* are paired in the same system; if *y* is *O1* then *y′* is *O2*). On each trial t, the posterior belief about *q_s_* is computed using Bayes rule, as follows:

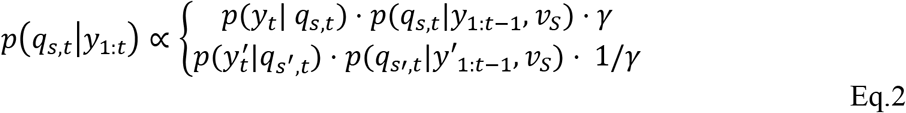

That is, *γ* = 1 for an ideal learner who can take advantage of the structural relationship 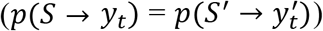 and learn from inferred outcomes as much as they learn from experienced outcome. Therefore, a participant with a higher level of *γ* is more likely learn from direct experiences (*S* → *y_t_*) but less likely to learn from inferred outcomes 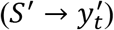. After each trial, the probabilities were normalized such that they remained bounded between 0 and 1.

While the likelihood *p*(*y_t_*|*q_s,t_*) or 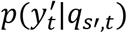 is *q_s,t_* or *q_s′,t_* respectively, we took into account the probability that the contingency of the system associated with the current choice (*S*) is reversed (*v_S_* = *p*(*J_s,t_* = 1)) when computing the prior, *p*(*q_s,t_*|*y*_1:*t*–1_). The term *v_S_* indicated the subjects’ belief that choice-outcome contingencies had reversed (*J_s,t_* = 1) for the chosen shape, *S*. Taken together, the prior belief of the associative contingency for a chosen shape remained the same as the previous trial (*p*(*q*_*s,t*–1_)) with the probability 1 – *v_S_* (if no reversal has occurred) or flipped to the inverse probability (1- *p*(*q*_*s,t*–1_)) with the probability *v_S_* (if a reversal has occurred). Therefore, the prior (*q_t_*|*y*_1:*t*–1_) is obtained by the following transition function:

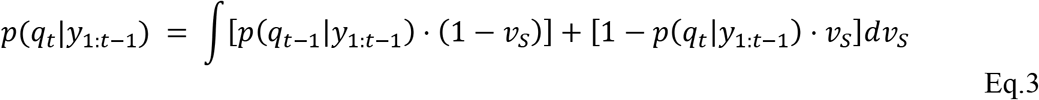

A second normalization step was done after applying the transition probabilities *v_s_* to the posterior probabilities of the current trial, such that the probability of all possible transitions equals to 1. Note that *v_S_* was defined and updated independently per four possible choice options. However, due to the inherent design of the underlying task structure, *v_S_* for shapes within the same system should be more correlated than *v_S_* of the other system. Finally, note that the reversal probability is fixed during the experiment but unknown to participants.

We then used the prior belief, in the associative contingencies, *p*(*q_s,t_*|*y*_1:*t*–1_), to compute the expected value of a given shape 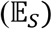 on each trial according to the following formula:

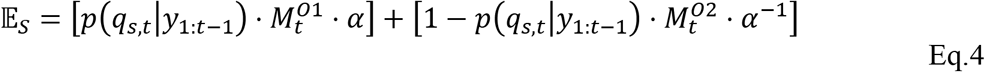

where *α* was a free parameter and reflected a subject’s preference for one outcome (*O1*) over the other (*O2*) (0 < *α* < ∞), and 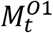 and 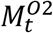 indicated the reward magnitudes of the outcome available in the current trial, *t*. We then predicted the choice of a participant between the two available shapes (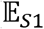 and 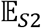) on each trial according to a SoftMax function:

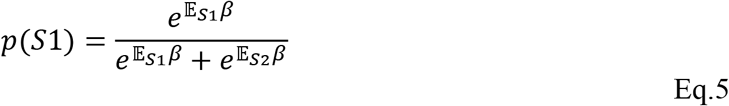

where the free parameter *β*, captured the level of sensitivity of choices to expected values (inverse temperature; 0 < *β* < ∞).

Finally, when the outcome was revealed, the reward prediction errors (rPE) were computed as follows:

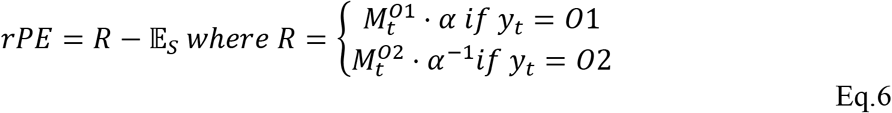

#### Alternative models

We tested the weighted-inference learning model against two additional models which made alternative assumptions about how subjects updated the posterior belief from the inferred outcomes. In the first alternative model, named the “perfect-inference model”, *γ* was fixed to 1 in Eq.2, resulting in equal and optimal integration for experienced and inferred outcomes (*γ_exp_*= *γ_inf_*=1). In the second alternative model, called the “no-inference model”, we assumed that participants did not take the structural relationship between shapes in the same system into the updates. Specifically, we set *γ_inf_* = 0 while *γ_exp_* = 1 in Eq.2. Therefore, an agent using no-inference model only learned from experienced outcomes but not from inferred outcomes.

#### Parameter estimates

The weighted-inference learning model has three free parameters, *α, β*, and *γ*, and two alternative models have two free parameters, *α*, and *β*. We fit all three models using custom Markov Chain Monte Carlo (MCMC) code in MATLAB R2018a. Each model was fit to maximize the likelihood of choices given model estimates of the expected value of each choice on each trial Eq.6.

#### Model Comparisons

To test potential overfitting, we compared the goodness of fit for each model type using the sum of the Bayesian Information Criterion (BIC) over subjects. This gave us an overall measure of how well the data were fit by each model at the group level, while penalizing models that added additional free parameters.

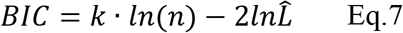

where *k* is the number of parameters in the learning model, *n* is the number of choices (i.e., trials) the subject made, and 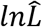 indicates the log-likelihood of each model.

#### Forward chaining cross-validation

We also tested if the weighted-inference model better predicts out-of-sample data. In the current study, a subject’s belief that any choice would lead to a specific outcome is dependent on the observations and inferences made in the preceding trials. That is the choice at the trial *t* cannot be predicted from any randomly sampled trials but only from *y*_1:*t*–1_. To account for the time-dependence of our data, we applied a forward chaining cross validation (CV) (Bergmeir & Benítez, 2012), which iteratively fits data from the earliest time points and uses the fitted model to predict later time points. We began by fitting the model on the first 20 trials of the experiment, then test the model on choices made in the 20 trials that came immediately after (trials 21 through 40). In the next iteration, we trained on the first 40 trials, and tested on choices made in the subsequent 20 trials (trials 41 through 60). This process continued in steps of 20 until the last iteration which trained on the first 140 trials and then were tested on the last 20 (total of 8 folds). We summed together the negative log-likelihood returned from each test set to determine which model performed best.

### Model free analysis of effects of decision history to the current decision

To test whether subjects showed a behavioral effect of learning on choice, we fit logistic regression models estimating the effects of past choice-outcome observations on which item was chosen at the current trial *t*. The regression model included the effect of experienced choice-outcome association going three trials back (denoted *t^E^* – *n*), and inferred choice-outcome relationships going three trials back (denoted *t^I^* – *n*), such as the following:

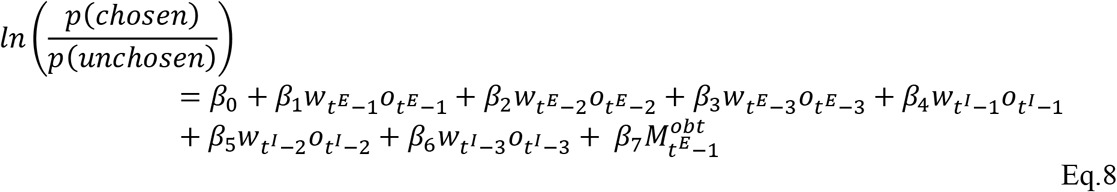

where *n* is the n-th previous trial that object was chosen, up to 3 previous experiences. For example, *t^E^* – 1 means the outcome directly experienced the last time they chose the current shape. The same notation is used for previously inferred outcomes. In this study, participants were presented two choice options among four shapes in each trial. This means that the chosen option in the current trial may not be available in the previous trial. As such, if current choice S or the paired shape, *S′* was not available in the previous trial, then *t^E^* – 1 or *t^I^* – 1 was the last trial when *S* or *S*′was chosen, respectively. We fit separate regression models for the choices of each of four shapes for each subject. For experienced trials, the value of each of these regressors was 1 if currently considered choice led to the desired outcome *n*-trials back and −1 if it did not. Thus:

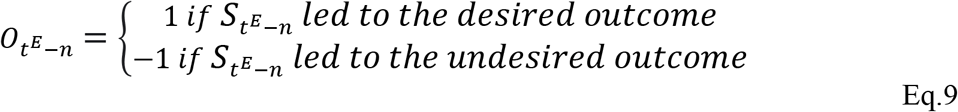

We also included contextual, counter-factual information about the other option in the experienced regressors. For example, if the subject were choosing between choices A and C but choose C and got the desired outcome, this may deter them from choosing shape A the next time A and C are available. We included this information for completeness with respect to all the experienced information that could influence the choice of a shape on any given trial.

For inferred trials, the regressor had a value of 1 *if the system pair* (i.e., B when participants’ choice is A in the current trial) led to the undesired outcome *n* trials back, such that

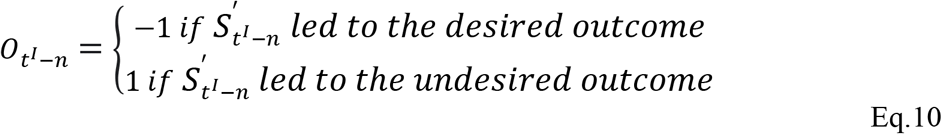

because this indicates that the currently considered shape should lead to the desired current.

We assumed that participants would desire the outcome with higher magnitude between *O1* and *O2*. To test the effects of greater desirability in previous choices in the current decision, we assigned the difference in reward magnitude 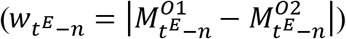 as a weight on each regressor. We did not consider the subjective preference of one outcome type over the other (*α* in the model, Eq.4) for the model free regression analysis. However, we repeated the analysis using *α* to moderate the value of each stimulus (Eq.4) to test if subjective preference produced any changes in these results. Finally, 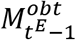 represents the influence of the magnitude of the reward obtained the last time subject chose the currently considered choice.

After fitting separate regression models for each fractal shape, we averaged together the regression coefficients (*β*) across shapes, representing the subject specific influence of previous decisions on the current choice.

### MRI data Acquisition

Data was acquired using Siemens Skyra 3 Tesla scanner. We used gradient-echo-planar imaging (EPI) pulse sequence, with a multi-band acceleration factor of 2, and set the slice angle of 30° relative to the anterior-posterior commissure line, minimizing the signal loss in the orbitofrontal cortex region (Weiskopf, Hutton, Josephs, & Deichmann, 2006). We acquired 38 axial slices, 3mm thick with the following parameters: repetition time (TR) = 1200 ms, echo time (TE) = 24 ms, flip angle = 67°, field of view (FoV) = 192mm, voxel size = 3 × 3 × 3 mm3. Contiguous slices were acquired in interleaved order. We also acquired a field map to correct for potential deformations with dual echo-time images covering the whole brain, with the following parameters: TR = 630 ms, TE1 = 10 ms, TE2 = 12.46 ms, flip angle = 40°, FoV = 192mm, voxel size = 3 × 3 × 3 mm3. For accurate registration of the EPIs to the standard space, we acquired a T1-weighted structural image using a magnetization-prepared rapid gradient echo sequence (MPRAGE) with the following parameters: TR = 1800 ms, TE = 2.96 ms, flip angle = 7°, FoV = 256mm, voxel size = 1 × 1 × 1 mm3.

### Preprocessing

Preprocessing of the data was done in *SPM12 (Wellcome Trust Centre for Neuroimaging*) in MATLAB (2018b Matworks). Data were preprocessed using the default options in SPM. Images were slice-time corrected and realigned to the first volume of each sequence. We realigned to correct for motion using a six-parameter rigid body transformation. Inhomogeneities in the field were corrected using the phase of non-EPI gradient echo images at 2 echo times, which were co-registered with structural maps. Images were then spatially normalized by warping subject specific images to the reference brain in the MNI (Montreal Neurological Institute) coordinate system with 2mm isotropic voxels. Finally, for the univariate analysis images were spatially smoothed using a gaussian kernel with full width at half maximum of 8mm.

### Univariate fMRI Analysis

To model BOLD activity in each voxel we used a GLM with four different regressors; the choice period (a boxcar, from the choice onset including the duration of .5s plus the reaction time of decisions), the button press (a stick function), the reward expectation period (a boxcar including jittered ISI) and the reward feedback phase (a 2 second boxcar). In the first GLM (GLM 1), we included the decision difficulty of each trial as a parametric regressor at the choice period. The decision difficulty was computed as the inverse of the expected value difference between options. See below:

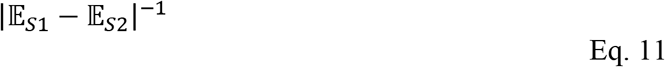

In addition, we computed the model-based belief updates to the choice-outcome associations after the outcome was observed in each trial and inputted this as a parametric regressor at the feedback phase. This belief update was calculated as the Kullback-Leibler divergence (*D_KL_*) between the prior and posterior belief in *q_s,t_* (Eq.1) for the chosen shape (*S*),

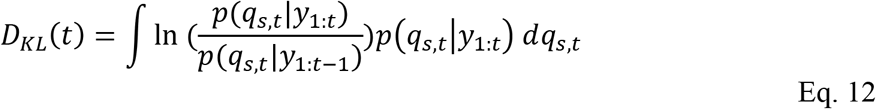

The *D_KL_* reflected changes in the model estimated “beliefs” about which choice led to which outcome (gift card identity) as participants progressed through learning. The *D_KL_* for shapes in each system were summed together to generate 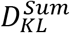. Six motion regressors were included as regressors of no interests in the model to account for translation and rotation in head position during the experiment. From the first-level analysis, contrast images of parameter estimates from regressors of the 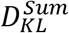 were estimated for each participant and inputted for the one sample t-test in the second level analysis.

We performed an additional GLM (GLM 2) to distinguish the neural activity reflecting the additional information gained from inference in belief updates at the time of feedback. To address this, we computed the *D_KL_* from the *no-inference model* 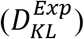 in addition to *D_KL_* which was estimated from the *weighted-inference model* given the subject specific weighting of inferred outcomes. We then generated 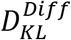 by subtracting 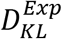 from the *D_KL_* of the *weighted-inference model* 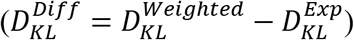. Thus, 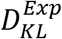 this would account for the update that comes from experiencing outcomes alone (i.e., no inference), whereas 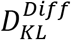 contained the additional updating that occurs when both inferred outcomes are integrated into a new belief. GLM2 was the same with the GLM1 except that we inputted two parametric regressors at the feedback phase.

### Group-level statistical inference

Group level testing was done using a one-sample t-test (df=36) on the cumulative functional maps generated by the first level analysis. All first level maps were smoothed prior to being combined and tested at the group level. To correct for multiple comparisons, we used Threshold-Free Cluster Enhancement (TFCE) which uses permutation testing and accounts for both the height and extent of the cluster (Smith & Nichols, 2009). All parameters were set to default parameters (H=2, E=0.5) and we used 5000 permutations for analysis. In all ROI based analyses and whole brain analyses we report effects that surpassed a p_TFCE_<.05 threshold.

We first performed group-level inference on independent anatomical ROIs, then performed exploratory whole brain analyses. For ROI analyses, we first extracted voxels from each ROI in each subject’s first-level activation map, averaged the maps together, then applied small volume TFCE correction. We used this analysis method for testing univariate effects of updating in VTA, decoding the latent cause of each system in mPFC and testing which regions represented association space. All other analyses were corrected for multiple comparisons at the whole brain level.

### Multivariate Analyses

The MPVA analysis aimed to identify regions of the brain that coded knowledge of the relationship between system pairs - the underlying structure of the task. To test this, we estimated the BOLD activity patterns during the feedback phase using unsmoothed preprocessed images. The feedback period was modeled as a boxcar that had a constant duration lasting 2 seconds from the feedback onset of each trial. No parametric modulators were added.

Each trial was labeled according to which shape was chosen and which outcome received from that choice (*S_t_ → y_t_*). Our main hypothesis of this study was that subjects would use knowledge about the underlying relationships between shapes in a system to make inferences of unobserved outcomes at feedback. If participants retrieve representations of these structural relationships at feedback to appropriately assign credit learned from experiences to the latent association, we could expect to decode the representations associated with the common causes that arise in trials where the systems’ pairs led to opposite outcomes 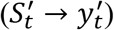. For example, trials where one shape in the pair lead to outcome one (e.g., A→*O1*) should share the same causes (e.g., *q*^1^) with trials where the other shape in the same system led to the opposite outcome (e.g., B→*O2*).

Importantly, to make sure that the activity patterns are not associated with the outcomes (*O1* or *O2*) presented on the screen but are associated with the latent causes (*q*^1^ or *q*^2^, i.e., the reward contingency in the system 1 and 2), we organized training and testing labels in a way to control for visual information. Specifically, we trained a shape against another shape which shared the same outcomes but did not share the causes (e.g., 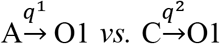) to identify the activity patterns specifically associated with the causes. Subsequently, we tested theses activity patterns on independent data sets which included the shapes that did not share the outcome with the training shapes but share the causes (e.g., 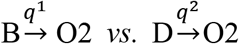). As this example showed, no sensory information was shared between training and testing sets that could influence the classifier to bias the results. See Table 1 for the full list of eight training and test pairs.

**Table 1:** Training and Testing Scheme of Linear Classifier for Latent Cause Decoding: This table shows all combinations of training left column) and testing (right column) trial sets used for decoding the latent cause at the time of feedback. Capital letters denote the chosen shape (A,B,C or D). Arrows followed by “O1” or “O2” indicate which outcome each shape led to on that trial. Note that training and test stimuli are matched for outcome identity so that no visual information can be used by the classifier to separate representations. Finally, letters above each arrow denote the latent cause (p or q) being decoded, indicating the system each stimulus belongs to (system 1 or system 2, respectively).

| Training Set | Test Set |
| --- | --- |
| $A \xrightarrow{q^1} O1$ vs. $C \xrightarrow{q^2} O1$ | $B \xrightarrow{q^1} O2$ vs. $D \xrightarrow{q^2} O2$ |
| $A \xrightarrow{q^1} O1$ vs. $D \xrightarrow{1-q^2} O1$ | $B \xrightarrow{q^1} O2$ vs. $C \xrightarrow{1-q^2} O2$ |
| $A \xrightarrow{1-q^1} O2$ vs. $C \xrightarrow{1-q^2} O2$ | $B \xrightarrow{1-q^1} O1$ vs. $D \xrightarrow{1-q^2} O1$ |
| $A \xrightarrow{1-q^1} O2$ vs. $D \xrightarrow{q^2} O2$ | $B \xrightarrow{1-q^1} O1$ vs. $C \xrightarrow{q^2} O1$ |
| $B \xrightarrow{1-q^1} O1$ vs. $C \xrightarrow{q^2} O1$ | $A \xrightarrow{1-q^1} O2$ vs. $D \xrightarrow{q^2} O2$ |
| $B \xrightarrow{1-q^1} O1$ vs. $D \xrightarrow{1-q^2} O1$ | $A \xrightarrow{1-q^1} O2$ vs. $C \xrightarrow{1-q^2} O2$ |
| $B \xrightarrow{q^1} O2$ vs. $C \xrightarrow{1-q^2} O2$ | $A \xrightarrow{q^1} O1$ vs. $D \xrightarrow{1-q^2} O1$ |
| $B \xrightarrow{q^1} O2$ vs. $D \xrightarrow{q^2} O2$ | $A \xrightarrow{q^1} O1$ vs. $C \xrightarrow{q^2} O1$ |

We then used a searchlight procedure to identify regions of the brain that contained representations of the underlying structure of the environment. Each searchlight consisted of a 5×5×5 voxel cube placed around a centroid voxel in the brain. Each centroid was required to values in at least 10 of the surrounding voxels to be considered for further processing and were then standardized by z-scoring the beta values within each searchlight.

The data were subset such that only the relevant trials were used for a particular classifier (see Table 1), then split by blocks into a training set and a test set. We used LIBSVM (Chang & Lin, 2011) to fit linear classifiers with training data, which we then used to classify data points from the test set. We iterated through this process for each of 2 blocks and for each of 8 combinations of training and test labels, then computed the mean decoding accuracy (average proportion of correct classifications) across all 16 classifiers for each voxel. The mean decoding accuracy for each voxel was compared to a voxel specific null distribution which was estimated with the same procedure while randomly assigning the labels over 100 permutations at each searchlight. The mean classification accuracy of this null distribution was subtracted off the classification accuracy of each searchlight to give us a measure of how reliably information about the latent cause could be decoded above chance. The resulting maps were then spatially smoothed using a gaussian kernel with full width half maximum of 8mm.

Group-level analyses were preformed using a one-sample t-test on accuracy maps across subjects (see Group-level Inference). For this analysis we focused on *a priori* defined ROIs in lOFC and mPFC (see selecting *a priori* ROIs) and corrected for multiple comparisons within each ROI using small volume correction TFCE. The threshold for significance remained the same (p_TFCE_ <.05)

### Trial-by-Trial Decoding Correlated with Belief Updates about Latent causes

To test whether the strength of the neural representations followed beliefs in specific choice-outcome contingencies, we correlated the probability that representations for the latent cause could be decoded on each trial with trial-by-trial belief updates in choice-outcome relationships. We used the same decoding procedure mentioned above to classify voxel patterns at feedback in each trial (see Multivariate Analyses), but additionally calculated the distance of each pattern from the hyperplane that divides categories. Distances were obtained using the equation specified on the LIBSVM webpage (https://www.csie.ntu.edu.tw/~cjlin/libsvm/faq.html). Patterns that are more distant from the hyperplane can be thought of as having more information about a category, and those that are closer to the hyperplane as having less information (Schuck & Niv, 2019). We then signed the distance of each point according to whether the predicted category label was correct (+ for correct, – for incorrect), and averaged the distance from each relevant decoding scheme. The signed distances were then regressed against the magnitude of the belief updates about the choice-outcome contingencies at feedback (*D_KL_*) estimated from the weighted-inference model.

The distances to the hyperplane and the magnitude of the *D_KL_* were then correlated using Spearman’s rank correlation, in each voxel throughout the brain. We used Spearman’s correlation as a conservative measure against outliers or nonlinear relationships that could bias the results. The correlation values were normalized using a Fisher transform and the resulting maps were spatially smoothed using a Gaussian kernel with full width at half maximum of 8mm. Group level analyses were preformed using a one-sample t-test on correlation values, then we applied TFCE correction to volumes within preselected ROIs. The same thresholds were applied for group level statistical correction (p_TFCE_ <.05).

### Representational similarity analysis (RSA) analysis of association space

We used RSA to look for regions of the brain that tracked the position of each system within an abstract association space as learning unfolded. If participants represented the state of each system as “positions” within an abstract association space, then we should observe similar neural representations when subjects occupy similar regions of the association space. For example, if subjects believe that the configuration of system 1 is A→*O1* and B→*O2* with probability *q*^1^=.80, the neural representation of this belief should be highly similar to a trial where participants believe system 1 is in the same configuration but *q*^1^=.75. However, the neural similarity should be more dissimilar if *q*^1^=.55, and yet more dissimilar if subjects believe that the configuration of system 1 has been reversed (A→*O2* and B → *O1*;*q*^1^ =.15). Note that while this example gives point estimates of *q*^1^, the true contingencies were defined as belief distributions which includes the confidence of each belief. Such increases in the dissimilarity of voxel patterns would suggest that neural representation is coded as an abstract value space, because it shows that distal points in the association space are represented with proportionately dissimilar activity patterns. As in previous work, we focused our analysis during the time of the ITI (Knudsen & Wallis, 2021).

To test this, we estimated the BOLD activity patterns during the ITI phase using unsmoothed preprocessed images. The ITI period was modeled as a boxcar and no parametric modulators were added. We created model representational dissimilarity matrices (RDMs) for each system 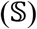 which measures the dissimilarity of seven factors of each trial (*t*) that could give rise to dissimilarity in neural representations. All RDMs were constructed such that they represented the dissimilarity of these factors between trials in separate blocks. The first two model RDM’s captured similarity of belief distributions across trials which were separated into the beliefs of the “task-relevant association” and “task-irrelevant association”. The task-relevant association RDM included the trial-by-trial dissimilarity between beliefs about 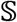. This included the trials in which participants used their belief about reward contingency to choose a particular shape and subsequently updated the belief with the given feedback. Therefore, the size of the task-relevant RDM corresponded to the number of trials in which a participant chose a shape associated with 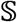 in block 1 × those trials in block 2. The task-irrelevant association RDM included the beliefs about reward contingencies for 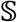, but it included the trials in which participants did not choose a shape associated with 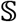, but needed to hold the representation for potential future or pending trials. Therefore, the size of the task-irrelevant RDM corresponded to the number of trials in which a participant chose a shape that was not associated with 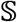 in block 1 × those trials in block 2. We computed the model RDMs of the task-relevant and -irrelevant contingencies in each of two systems.

To compute the trial-by-trial dissimilarity between two belief distributions across sessions, we used the Jensen-Shannon Divergence (*D_JS_*) between distributions. This metric is commonly used to measure the dissimilarity between two distributions (*D*_1_ *and D*_2_). Note that *D_JS_* is symmetric. That is, *D_JS_*(*D*_1_||*D*_2_) is the same with *D_JS_*(*D*_2_||*D*_1_) unlike the KL divergence. We computed *D_JS_* by combining *D_KL_* of each distribution to their mean distribution 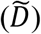:

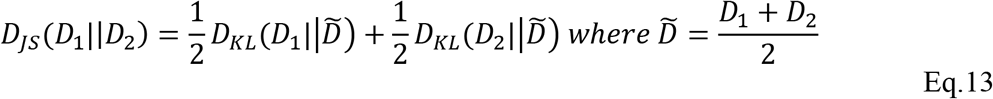

We included 5 additional model RDM’s to control for alternative possible explanations of neural similarity. These were as follows: the identity of the chosen shape (CS), the identity of the unchosen shape (US), choice location (right or left side; CL), the outcome identity (OI), and the signed reward prediction error (rPE) computed by the weighted-inference model (Eq.7). All of these model RDMs were binary matrices except for the rPE matrix, in which the dissimilarity was computed as the absolute difference in rPE’s between trials. All analyses were conducted separately per system (see Fig.S6 for correlation matrix).

We then followed the same searchlight procedure as with the MVPA (5×5×5 voxel cube around a centroid voxel), at each centroid generated a neural RDM by calculating the Euclidean distance between voxel activities for trials in each session, after standardizing voxel activity within the ROI. We then regressed the neural similarity matrix against each of the model RDM’s (Flesch et al., 2021; Parkinson et al., 2017) using the following GLM:

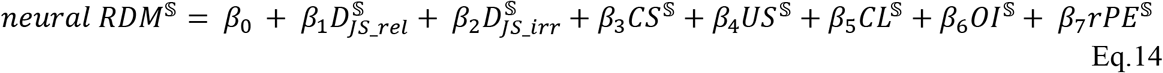

All predictors were z-scored before fitting the GLM. Each subject’s resulting 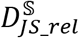 beta maps were averaged across systems to produce a single estimate of the correlation between system specific belief similarity and neural similarity. Group level analyses were preformed using a one-sample t-test on smoothed beta values. We applied TFCE correction to volumes within preselected ROIs at p_TFCE_ <.05 threshold for group level statistical correction.

To visual this analysis and create Fig.4C, we performed the following procedure: First, we split the data into a left-out run and a visualized run. For the left-out run, we created two “template patterns” that represented the average multivariate pattern for each of the two possible configurations for a given system. For example, we created a template pattern for the state of system 1 when A→*O1* and B→*O2* ( *q*^1^) by averaging the activity in each voxel across trials in which the weighted-inference model indicated this belief was true. The same was done for all other possible configurations of each system (1 – *q*^1^, *q*^2^, 1 – *q*^2^). These template patterns were then used as a comparison for trials in the visualized block, pre- and post-reversal. Reversal points were identified as trials that subjects had a different beliefs about the configuration of a system compared to the trial before it (e.g., *q*^1^ flipped to 1 – *q*^1^). All reversal points were required to have at least 3 prior trials in which the same belief was held by the learning model (e.g., *q*^1^) and three trials after when the configuration changed (1 – *q*^1^). We then compared these trials to the template pattern that matched the belief prior to the reversal, such that if prior to the reversal the learner’s belief was that *q*^1^ was true, the neural pattern of those trials was compared to the template pattern for *q*^1^. Similarity patterns were compared using spearman’s rank correlations. However, no statistical inference was conducted on the correlations, as they were only used to visualize the analysis conducted in figure 4B.

### Selecting *a priori* ROIs

Regions of interest in prefrontal cortex were generated from anatomically defined regions in the Human Connectome Project Dataset (Glasser et al., 2016). The OFC ROIs corresponded to bilateral area BA13 (index 92) and for the mPFC we used BA10 (index 65). We included these regions because they have been previously implicated in credit assignment for causal choices, particularly in similar contingency learning tasks (Boorman et al., 2013, 2016; Jocham et al., 2016). To understand the role of dopaminergic regions of the midbrain in inferential updating, we looked at the ventral tegmental area (VTA) which has previously been linked to updating choice-outcome association (Boorman et al., 2016; Gershman & Uchida, 2019; Howard & Kahnt, 2018; Iglesias et al., 2013). Here, we used an anatomical VTA\SN ROI taken from a previous study linking the VTA to social updating about the trustworthiness of advice from others (Diaconescu et al., 2017).

## Supplemental Information

**Fig.S1.**
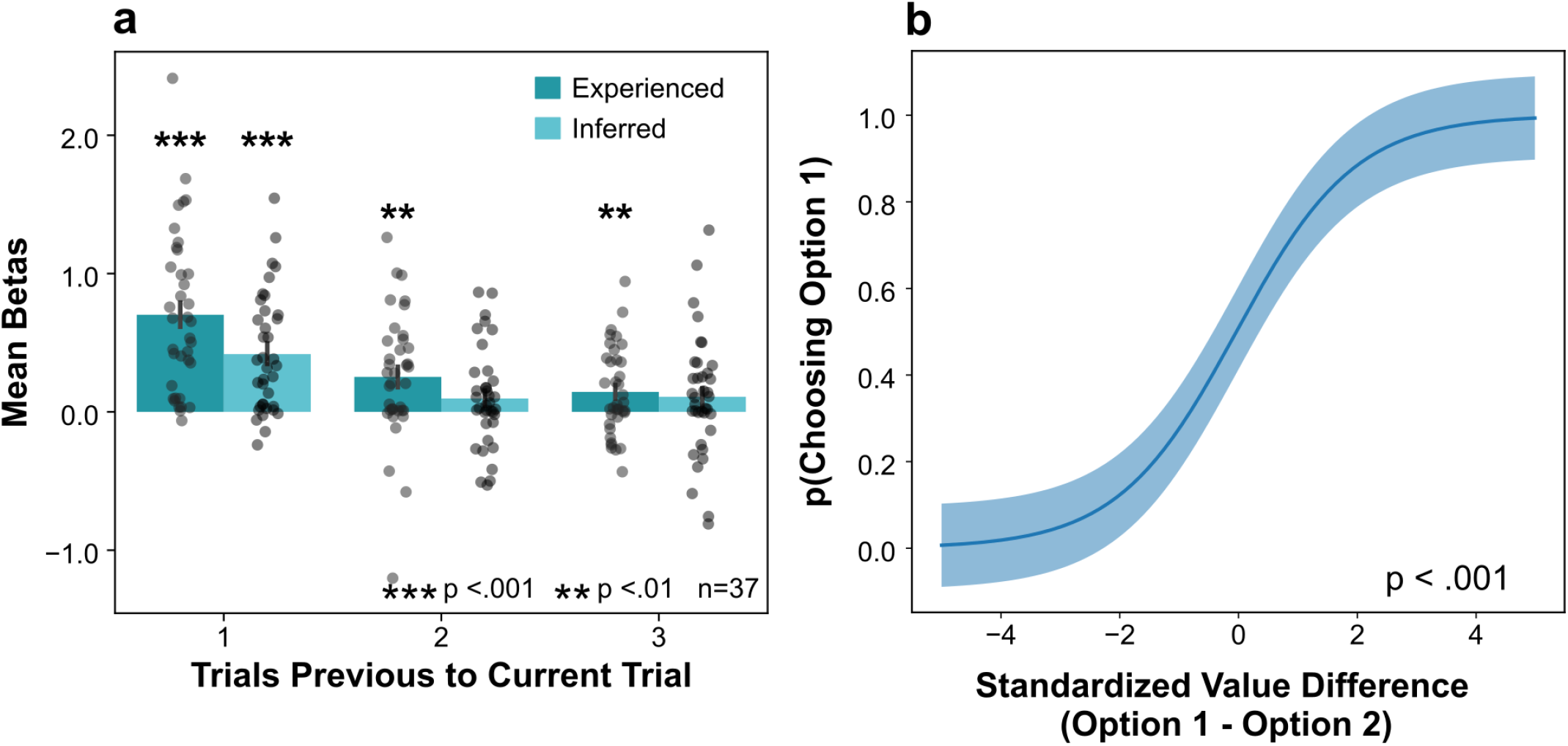
Behavioral Results Including Preference Term. **a)** Results of a logistic regression analysis which shows the influence of past choices and outcomes on the current choice. This model includes a preference term (*α*) for each outcome identity when computing the value if each possible outcome in the current trial. Both experienced and inferred past choice-outcome associations significantly predicted current choice. Height of the bars represents the mean of regression coefficients ± SEM. As expected, this influence decreased for trials further in the past. **b)** Logistic regression model predicting the probability of choosing Option 1 given the standardized (z-scored) difference in value for each option.

**Fig.S2.**
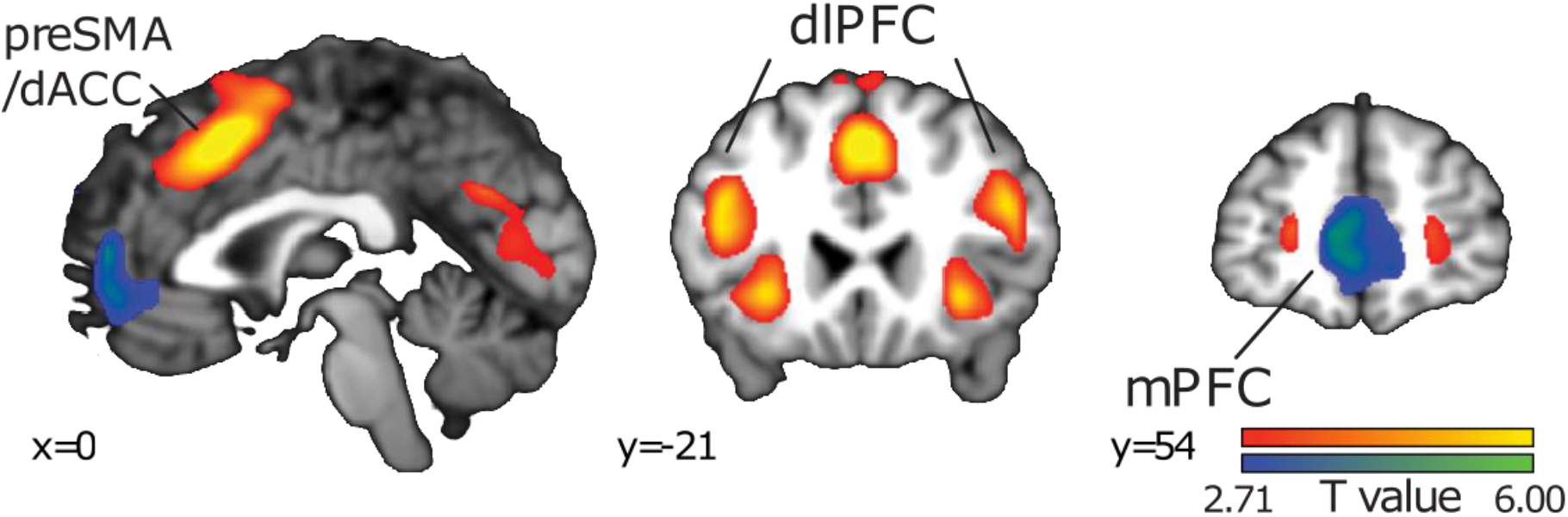
Network of Regions that Reflect Updating Signals Combining Experienced and Inferred Information. Sagittal and coronal slices showing t-statistic maps that display neural regions reflecting the updating of the stimulus outcome contingencies by integrating both inferred and experienced outcomes 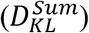. Maps display regions at a threshold of t(36)=2.71, p<.005).

**Fig.S3.**
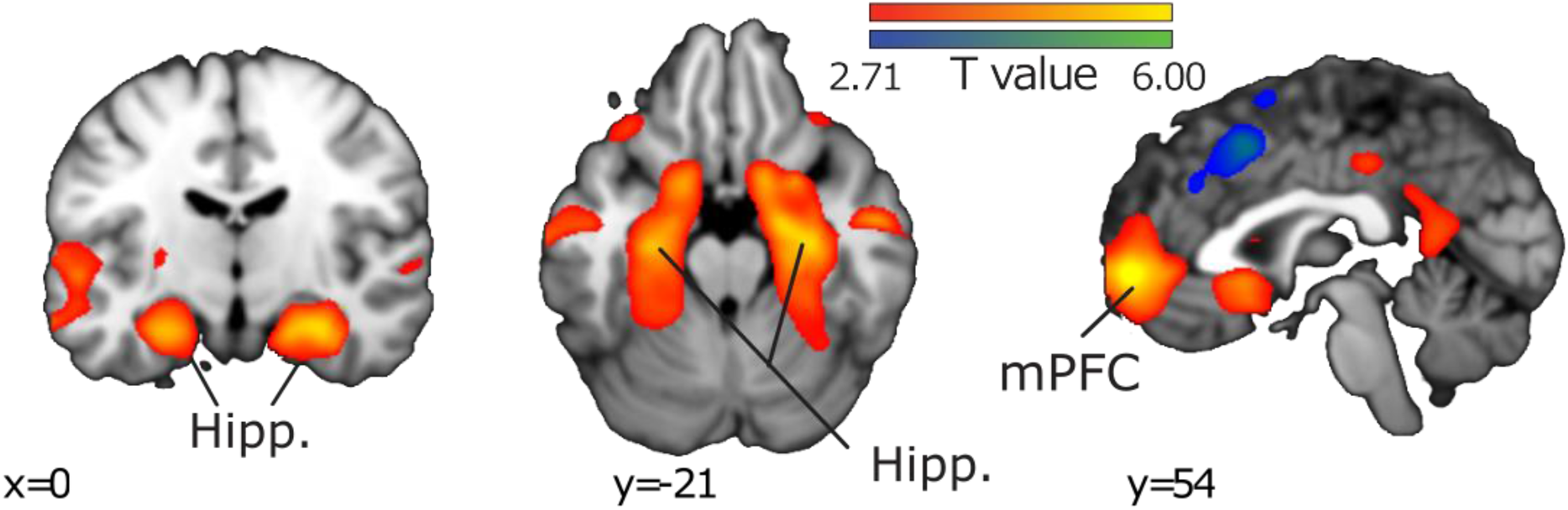
Network of Regions Whose Activity Reflects Reward Prediction Error. Regions encoding the reward prediction error (*RPE*) at the time of feedback (EQ.7). Maps are displayed with the same conventions as in Fig.S2. Clusters in Hippocampus and mPFC survived TFCE correction at the whole brain level (p_TFCE_ <.05). Note the absence of any effect in VTA.

**Fig.S4.**
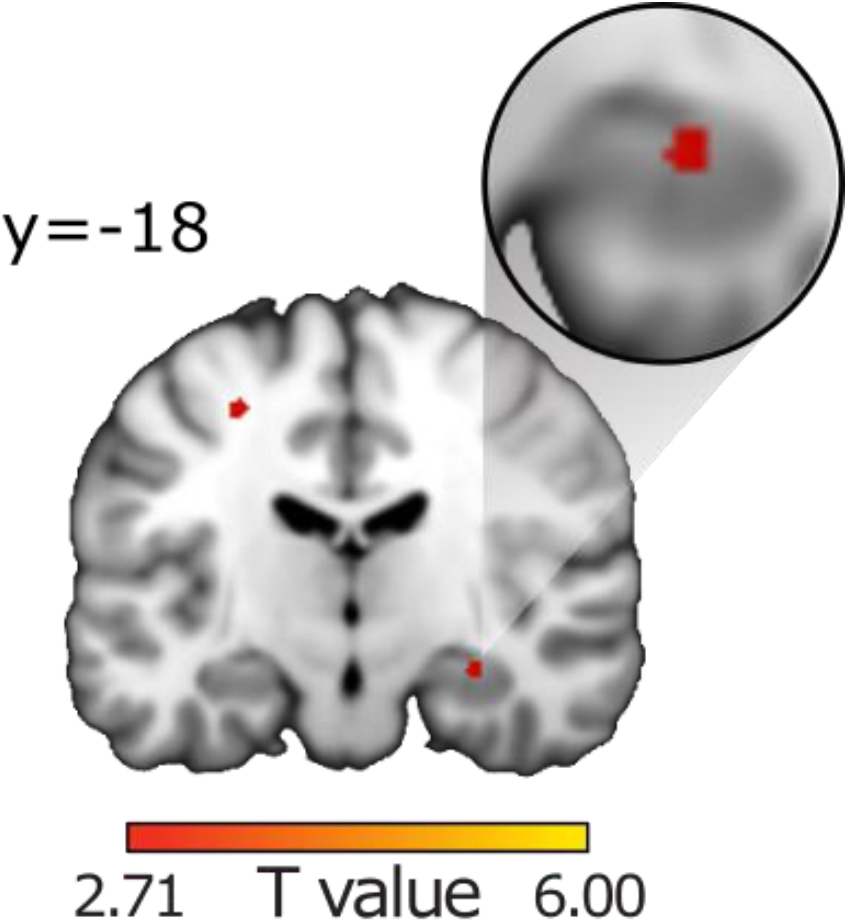
Hippocampal Tracking of Inferred Position. Coronal slice through t-statistic map showing a hippocampal region in which the model RDM was positively related to the neural RDM. Maps are displayed with the same conventions as in Fig.S2.

**Fig.S5.**
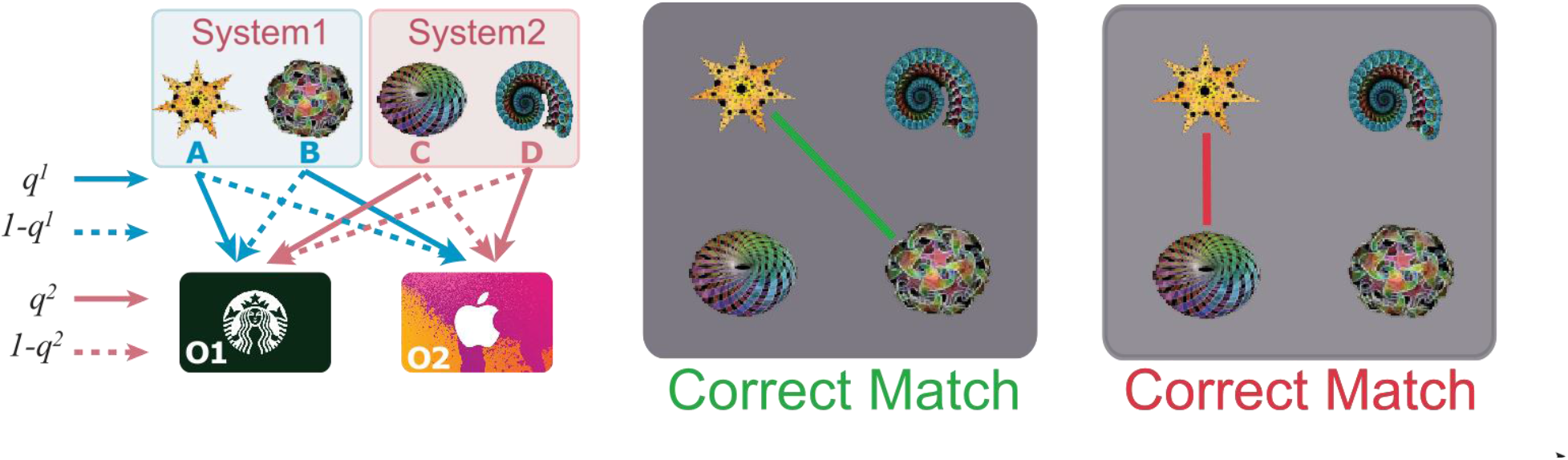
Matching Task. Matching task completed by subject during learning of the causal inverse relationships. Items were randomly displayed in 4 quadrants of the screen, and participants were asked to match items of the same systems by consecutively pressing the numbers associated with each quadrant. For example, subjects would match the top left and bottom right corners by pressing 2 and 4 consecutively. Correct matches were indicated by a green line connecting the shapes and incorrect responses were indicated by connecting shapes with a red line.

**Fig.S6.**
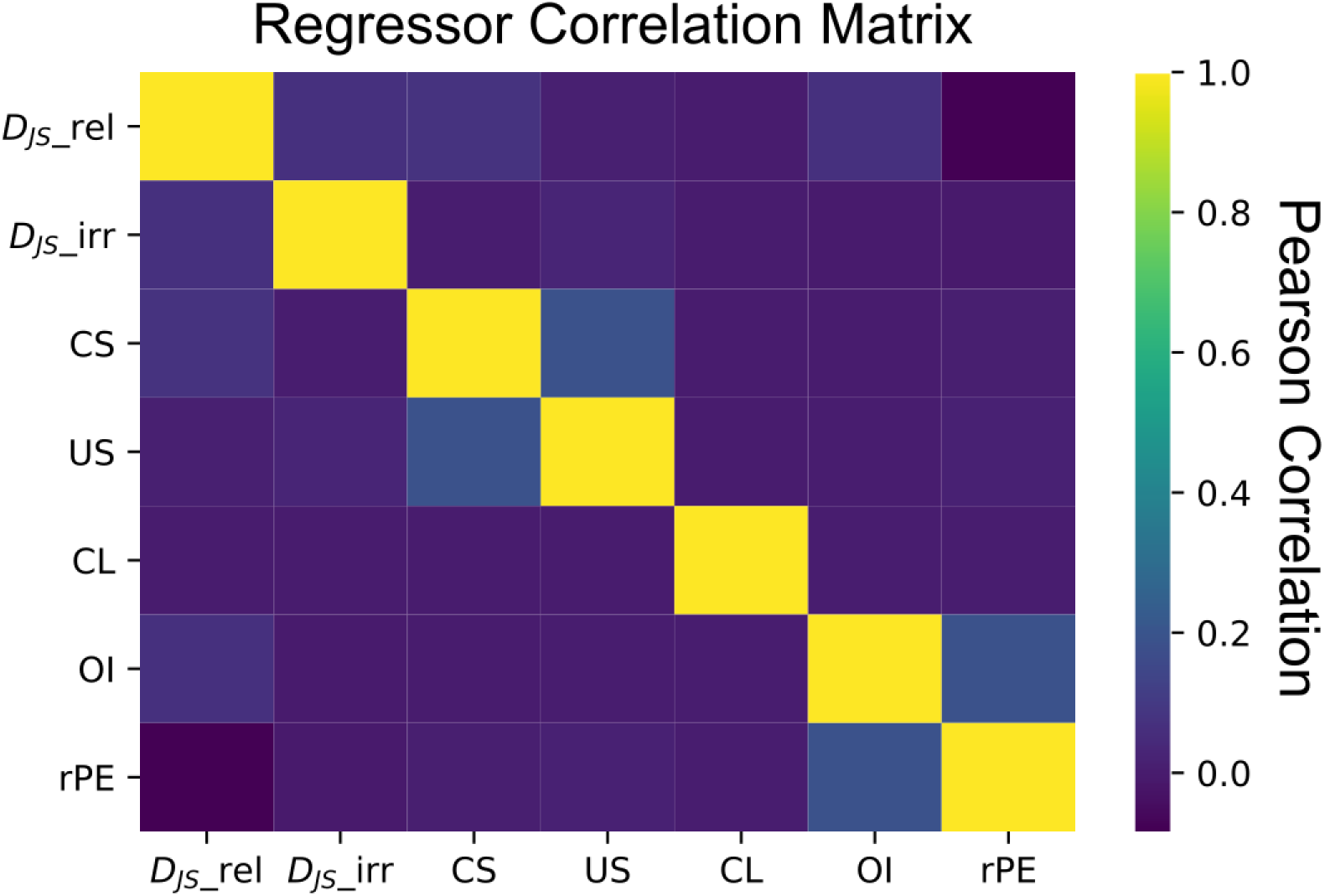
Correlation Matrix of Regressors in Position Tracking Analysis. Average Pearson correlations for all predictor variables in the position tracking analysis (Fig.4). All correlations were averaged over subjects and systems. The regressors were as follows: the *D_JS_* of the currently updated (relevant) system (*D_JS__*rel), *D_JS_* of the irrelevant system (*D_JS__*irr), the identity of the chosen shape (CS), the identity of the unchosen shape (US), choice location (right or left side; CL), the outcome identity (OI), and the signed reward prediction error (rPE) computed by the weighted-inference model.

**Table S1.** Distributions of parameter estimates for each behavioral model. fMRI activation table

| *Model* | $\alpha$ Median | $\alpha$ IQR | $\beta$ Median | $\beta$ SD | $\gamma$ Median | $\gamma$ SD |
| --- | --- | --- | --- | --- | --- | --- |
| *Weighted Inference* | 1.04 | 0.49 | 0.04 | 0.045 | 1.29 | 1.27 |
| *No Inference* | .99 | 0.44 | .049 | 0.05 | 0 | 0 |
| *Perfect Inference* | 1.03 | .56 | .032 | .033 | 1 | 0 |

**Table S2.** Univariate Activation at the Time of Feedback Given 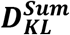 Update (related to Fig.S2): ✶ indicates TFCE correction with an independently defined ROI.

| *Region* | *cluster size (k)* | *p-value (uncorrected)* | *p-value TFCE* | *Peak (MNI)* |  |  |  |
| --- | --- | --- | --- | --- | --- | --- | --- |
|  |  |  |  | *t-stat* | *x* | *y* | *z* |
| L.Ant. Insula | 210 | 2.78x10<sup>-6</sup> | .005 | 5.69 | -30 | 22 | 2 |
| R.Ant. Insula | 93 | 7.90x10<sup>-6</sup> | .02 | 5.35 | 32 | 26 | 0 |
| L. Dorsolateral PFC | 1209 | $5.42 \times 10^{-7}$ | $4.00 \times 10^{-4}$ | 6.22 | -36 | 8 | 36 |
| R. Dorsolateral PFC | 707 | $1.45 \times 10^{-6}$ | .002 | 5.9 | 46 | 24 | 28 |
| preSMA/Anterior Cingulate Cortex | 1319 | $1.93 \times 10^{-8}$ | 0 | 7.31 | 0 | 18 | 50 |
| mPFC | 310 | $3.61 \times 10^{-5}$ | .0720 | 4.85 | -4 | 56 | 12 |

**Table S3.** Univariate Activation at the Time of Feedback Given 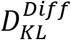 Difference Magnitude (related to Fig. 2): ✶ indicates TFCE correction with an anatomically independent defined ROI.

|  |  |  |  | Peak (MNI) |  |  |  |
| --- | --- | --- | --- | --- | --- | --- | --- |
| Region | Voxels | p-value | p-value TFCE | t-stat (max) | x | y | z |
| L. Anterior Insula | 116 | $7.38 \times 10^{-7}$ | .028 | 6.12 | -30 | 22 | -4 |
| R. Anterior Insula | 150 | $1.21 \times 10^{-6}$ | .029 | 5.58 | 32 | 22 | -2 |
| L. Dorsolateral PFC | 649 | $1.18 \times 10^{-5}$ | .025 | 5.22 | -44 | 24 | 28 |
| R. Dorsolateral PFC | 617 | $1.97 \times 10^{-5}$ | .025 | 5.05 | 44 | 26 | 28 |
| Pre-Supplementary Motor Area/ Anterior Cingulate Cortex | 557 | $1.86 \times 10^{-6}$ | .015 | 5.86 | 4 | 20 | 50 |
| Ventral Tegmental Area\Substantia Nigra★ | 89 | .0005 | .01 | 3.99 | 2 | -18 | -10 |

**Table S4.** Decoding of the latent Cause Magnitude (related to Fig. 3B): ✶ indicates TFCE correction with anatomically independent defined ROI

|  |  |  |  | Peak (MNI) |  |  |  |
| --- | --- | --- | --- | --- | --- | --- | --- |
| Region | Voxels | p-value (uncorrected) | p-value TFCE | t-stat (max) | x | y | z |
| mPFC★ | 141 | .002 | .008 | 3.54 | -6 | 50 | -10 |

**Table S5.** Decodability of Latent Cause Correlated with Model Based Update (related to Fig. 3C): ✶ indicates TFCE correction with an anatomically independent defined ROI

|  |  |  |  | Peak<br>(MNI) |  |  |  |
| --- | --- | --- | --- | --- | --- | --- | --- |
| Region | Voxels | p-value | p-value<br>TFCE | t-stat<br>(max) | x | y | z |
| mPFC ★ | 44 | 2.55x10<sup>-4</sup> | .0053 | 4.19 | 8 | 46 | -10 |
| Ventral<br>Tegmental<br>Area\Substantia<br>Nigra ★ | 63 | 7.30<sup>-4</sup> | .014 | 3.82 | -2 | -20 | -12 |

**Table S6.** Tracking Position of Systems within Association Space (related to Fig. 4): ✶ indicates TFCE correction with an anatomically independent defined ROI

|  |  |  |  | Peak<br>(MNI) |  |  |  |
| --- | --- | --- | --- | --- | --- | --- | --- |
| Region | Voxels | p-value | p-value<br>TFCE | t-stat<br>(max) | x | y | z |
| L.<br>Posterior<br>OFC ★ | 25 | .0007 | .002 | 4.24 | -26 | 30 | -12 |
| R.<br>Posterior<br>OFC ★ | 23 | .0002 | .007 | 3.78 | 28 | 28 | -14 |
| mPFC★ | 94 | 8.61x10<sup>-5</sup> | .0008 | 4.56 | 4 | 58 | -4 |
| Vis.<br>Cortex | 10602 | 4.14x10<sup>-6</sup> | 2.71x10<sup>-17</sup> | 5.65 | 6 | -72 | 2 |

